# Atomistic modeling towards predictive cardiotoxicity

**DOI:** 10.1101/635441

**Authors:** Kevin R. DeMarco, John R. D. Dawson, Pei-Chi Yang, Slava Bekker, Van A. Ngo, Sergei Y. Noskov, Vladimir Yarov-Yarovoy, Colleen E. Clancy, Igor Vorobyov

**Affiliations:** Department of Physiology and Membrane Biology, University of California, Davis, CA, USA; Department of Pharmacology, University of California, Davis, CA, USA; Biophysics Graduate Group, University of California Davis, CA, USA; American River College, Sacramento, CA, USA; Centre for Molecular Simulation, Department of Biological Sciences, University of Calgary, Calgary, Alberta, Canada

**Author notes:** Correspondence: Igor Vorobyov, Ph.D. Department of Physiology and Membrane Biology, University of California, Davis, Tupper Hall, RM 4133, Davis, CA 95616-8636. Colleen E. Clancy, Ph.D. Department of Physiology and Membrane Biology, University of California, Davis, Tupper Hall, RM 4303, Davis, CA 95616-8636.

**Keywords:** hERG, dofetilide, safety pharmacology, molecular modeling, molecular dynamics

## Abstract

Current methods for assessing safety pharmacology in the context of cardiac arrhythmia risk are unable to distinguish between drugs that cause cardiac rhythm disturbances and benign drugs. Drugs deemed likely to be unsafe share the common property of blocking the human Ether-à-go-go-Related Gene (hERG) encoded cardiac potassium channel and consequent prolongation of QT interval on the ECG. However, hERG block and QT prolongation alone are not selective indicators for cardiac arrhythmia. Here we present a prototype computational framework to distinguish between safe and unsafe hERG blockers. We used recent cryo-EM hERG structure to build and validate an atomistic structural model of the channel open conducting state. We also developed structural atomistic models of dofetilide, a hERG blocking drug with high pro-arrhythmia risk, in both charged and neutral ionization states. Next, we employed unbiased and enhanced sampling all-atom molecular dynamics (MD) simulations to probe atomic-scale mechanisms of dofetilide interaction with open-state hERG. Multi-microsecond drug “flooding” simulations revealed spontaneous dofetilide binding to the channel pore through the intracellular gate. Umbrella sampling MD was used to compute dofetilide affinity to hERG, in good agreement with experiment, as well as ingress and egress rates, which in a novel linkage between the atomistic and functional scale are utilized in our companion paper (Yang P-C *et al.* 2019 *bioRxiv*:635433) to parameterize functional kinetic models of dofetilide - hERG interactions used to predict emergent drug effects on the cardiac rhythm. This study represents the first necessary components of a computational framework for virtual cardiac safety pharmacology screening from the atom to the rhythm.

## Introduction

*“Poisons and medicine are often the same substance given with different intents*” wrote Peter Mere Latham in the 19th-century (1). Drug binding to a receptor protein is determined by the chemical composition of the drug and its interaction with the local protein environment (2). Even slight modification to drug chemistry may impact the potency of binding, and the resulting functional impact can vary widely from therapeutic to lethal (3). Although these principles are well understood (4), it has been an exceedingly difficult problem to predict the link between drug chemistry and altered physiological function. A persistent example of drug failure that cannot be predicted from drug chemical composition is unanticipated cardiotoxicity in the form of deadly abnormal rhythms in patients resulting from block of the major repolarizing potassium channel hERG (5).

Unpredictable drug-induced arrhythmia has been estimated to affect pharmaceuticals from multiple drug classes (6), with an estimated 3% of all prescription drugs worldwide harboring pro-arrhythmic side effects (7). The problem has even plagued drugs intended to treat arrhythmias, with drugs from nearly all antiarrhythmic drug classes being linked to drug induced arrhythmia (8, 9). Indeed, cardiotoxicity is one of the leading causes of drug attrition (10), and accounts for 22-28% of US post-marketing drug withdrawal (11).

Experimental studies alone cannot provide the necessary detailed structural information on drug interaction with ion channels. For instance, liquid chromatography with mass spectrometry can elucidate pharmacokinetic information (12, 13), mutagenesis studies and voltage patch-clamp techniques can tentatively identify specific residues involved in drug – channel interactions (14–16). Experimental techniques for determining high-resolution structures of drug-channel complexes at the atomic scale have their own limitations such as system size (NMR), bound state stability (X-ray) and resolution (cryogenic electron microscopy (cryo-EM) (17, 18). The most important limitation for understanding drug-protein interactions, however, is the static nature of information provided by a single reported structure. It is well-established that the dynamic adaptation of the binding pocket in a protein receptor with bound drug, and protein dynamics itself, are central for understanding high-affinity and specificity of protein-ligand recognition (19). Another important limitation in using a single receptor structure determined by NMR, X-Ray or cryo-EM techniques is related to an apparent need to sample all relevant conformational states for substrate and receptor itself and assigning their Boltzmann weights accurately (20).

The availability of high-resolution structures of ion channels and robust computational modeling tools now make it possible to predict the impact of a drug molecule on the emergent electrical activity in the heart. Indeed, the structures of multiple ion channels at atomistic resolution have been determined in recent years via X-ray crystallography and cryo-EM (21, 22). Significantly, a structure of the hERG channel in a putative open state was recently determined at a 3.8 Å resolution by cryo-EM (23) and was used in this study. The advances in molecular modeling and simulation methodology (24) and availability of powerful and efficient high-performance computing resources (25, 26) have made rigorous atomic-resolution investigation of ion channel function (27), and ion channel – drug interactions (28) now feasible.

In this study, we show that it is possible to predict drug-channel kinetics from atomic-resolution simulations, and hence begin to solve problem of preclinical drug cardiac safety screening *in silico.* Our current work expands upon our recent investigations (29–35) into drug and ion channel model development, and drug-channel binding using all-atom MD simulations. For instance, we recently developed and validated atomistic force field models of charged and neutral forms of the local anesthetic and anti-arrhythmic drug lidocaine and determined its preferred binding modalities and pathways in the cardiac voltage-gated sodium (Na_V_) channel using multi-microsecond long unbiased MD simulations, providing good agreement with available experimental electrophysiology data as well as new structural predictions (35). In another recent work, we developed force field models of the hERG-blocking drug sotalol, which has high pro-arrhythmia proclivity, and studied its lipid membrane partitioning using both unbiased and enhanced sampling MD simulations, obtaining reasonably accurate energetics and kinetics of drug – lipid bilayer interactions (34).

To develop our multi-scale prototype methodology, we focus here on another potent hERG blocker dofetilide (36), which has a high proarrhythmic risk and has been extensively studied experimentally (37–39). Many drugs like dofetilide and sotalol are weak bases, and therefore exist in equilibrium between ionized and neutral fractions in aqueous solution at physiological pH of 7.2. Both ionized and neutral drug forms are capable of accessing the hERG channel pore through an intracellular aqueous, but the neutral form is potentially capable of interacting with the channel by passage through lipid-facing fenestrations, as was proposed over 40 years ago (40), suggested by recent channel structures (41, 42) and supported by our local anesthetic – Na_V_ channel atomistic simulations (30, 35).

In an earlier study (33), we used preliminary dofetilide force field models and homology models of the hERG pore domain, based on then-available K_V_1.2/2.1 and KcsA structures (29), to study effect of pH on the channel – drug interactions. We have recently integrated our atomistic simulation results using these models into multi-scale functional modeling of hormone effects on arrhythmogenesis induced by hERG mutations or drug channel block (32). In another study, we used MD simulation results to gain insight into possible molecular mechanisms of flecainide action in treatment of catecholaminergic polymorphic ventricular tachycardia (CPVT) via multi-scale functional modeling (31). However, in both cases such integration was limited by *qualitative* information about drug – lipid membrane (31) or drug / hormone – channel interactions (32), which was not directly used to parameterize functional models.

Furthermore, the published structure of the putatively open-state for the hERG channel (23) has challenged many of the previously-proposed modes of drug binding. The commonly accepted idea emphasized the importance of a wide and easily accessible intra-cavitary site provided a combination of amphipathic-to-polar (pore-helix residues) and hydrophobic-aromatic moieties (S6 helix). Therefore, the intra-cavitary binding site provides a high-affinity/low-specificity binding pocket targeted by a broad panel of pharmaceutics ranging from anti-inflammatory to anti-convulsants and anti-arrhythmic drugs (43, 44). In striking contrast, the high-resolution cryo-EM structure of the hERG intra-cellular cavity shows a very constricted space with an obvious hydrophobic character defined by the positions of S6 helix Y652 and F656 residues (23). Here, we utilized this hERG channel structure to develop an atomic-scale model of hERG in an open conducting state, supported by our multi-microsecond-long MD simulations under applied voltage. Optimized structural atomistic models of charged and neutral dofetilide, compatible with biomolecular CHARMM force fields (45), were developed and validated using lipid membrane partitioning molecular dynamics (MD) simulations. Then, we used multi-microsecond-long unbiased MD simulations to identify potential drug binding sites, entry and egress pathways. Finally, we used biased, one-dimensional umbrella sampling (US) MD simulations (46) to estimate binding affinities and association/dissociation rates (k_on_/k_off_) of charged and neutral dofetilide with open state hERG. These data are then used to inform functional scale models for predicting drug pro-arrhythmia risks, as described in the companion paper (47).

The combination of computational modeling and simulation tools used in this study was optimized to allow for robust and accurate development and validation of atomistic *in silico* models of the channel-drug interactions. This approach allowed us to obtain quantitative estimates of state-dependent drug – channel binding affinities and rates, and then to *directly* utilize these quantities as parameters in our functional kinetic model of hERG – dofetilide interactions, as described in the companion paper (47). This provides a novel link between our atomistic and functional kinetic models of *in silico* safety pharmacology and allows for prediction of a drug pro-arrhythmia proclivity based on its chemical structure – all the way from the atom to the rhythm.

## Results

The goal of this study was to develop an accurate and robust quantitative atomic-scale model of the hERG channel – dofetilide interactions for a particular channel conformational state taking into account different drug ionization states and their channel binding pathways, as outlined in **Figure 1A**. To accomplish this goal, we developed and validated atomistic force field models of dofetilide in both charged and neutral ionization states (red and green hexagons in **Figure 1A**). We then developed and validated an atomistic structural model of the hERG channel in an open conducting state based on the recent cryo-EM structure, shown schematically in **Figure 1A**. Next, we employed a variety of unbiased and enhanced sampling all-atom MD techniques to probe atomic scale mechanisms of dofetilide interaction with the hERG open channel. We utilized computed energetic and dynamic quantities to determine dofetilide ingress and egress rates to be used as parameters in functional protein-scale channel – drug model (outlined in **Figure 1B**) and subsequent cell- and tissue-scale simulations discussed in the companion paper (47).

**Figure 1.**
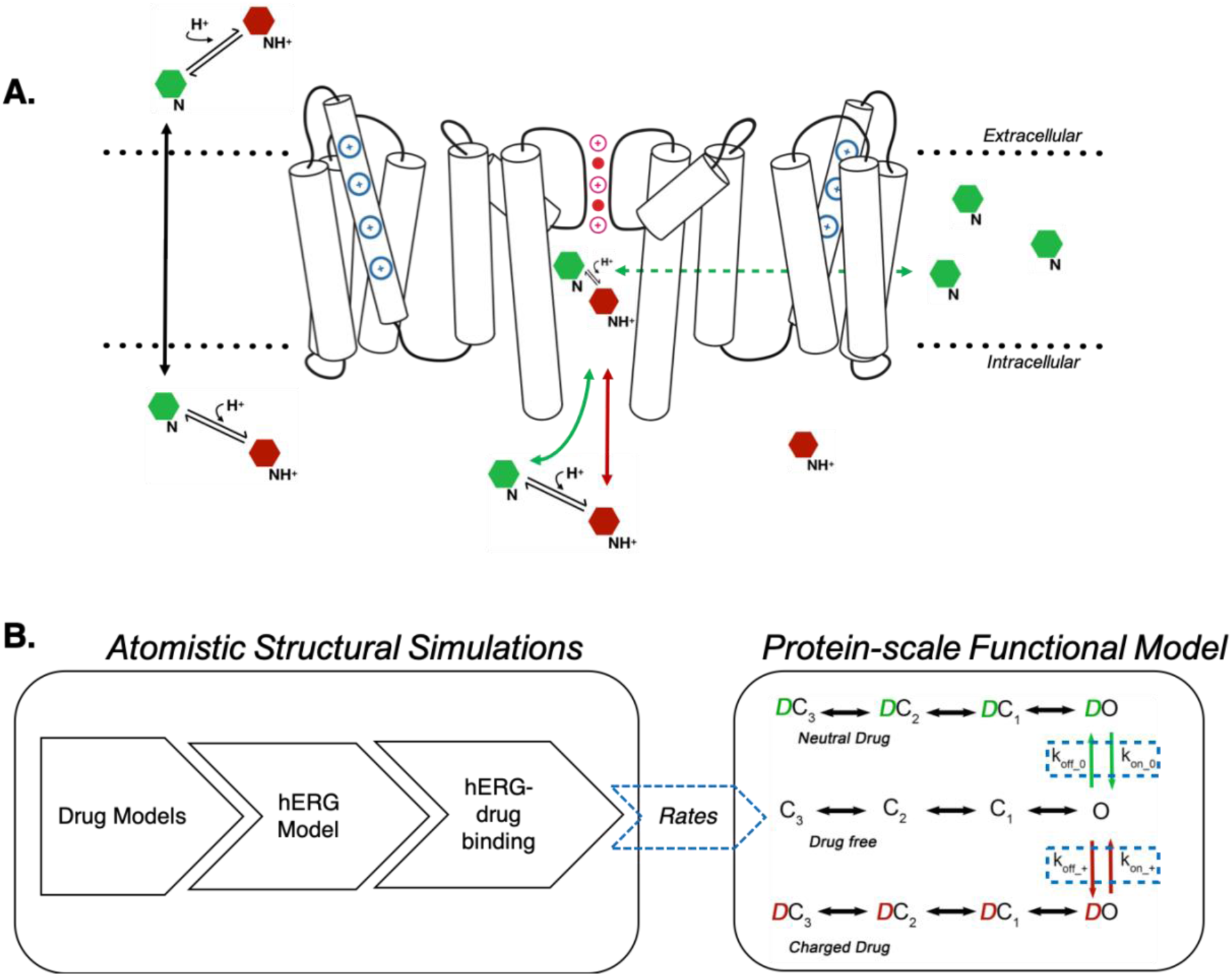
Overview of simulated system topology and multi-scale pipeline. **(A)** Structural illustration of a simulation system with the channel, lipid membrane and drug molecules shown. An open conducting hERG potassium channel is embedded in a lipid membrane with a single file of alternating K^+^ ions (purple) and water molecules (red) in the selectivity filter, and voltage sensors in the activated position, with α-helices depicted as white cylinders. Drug molecules are depicted as hexagons in equilibrium between charged (red) and neutral states (green). Drug binding pathways are depicted by red of green arrows. **(B)** Flow chart diagram depicting the topology of our multi-scale modeling pipeline, going from atom-scale studies (left) to computing rates that are used in channel Markov state model, subsequently utilized for cell and tissue simulations in the companion paper (47) (right).

### Dofetilide Models and Membrane Interactions

Dofetilide has a basic (protonatable) tertiary amine group and two acidic sulfonamide groups with estimated aqueous p*K*_a_ values of of 7.0, 9.0 and 9.6, respectively,(48) and therefore at physiological pH of 7.2 exists in equilibrium between neutral (Dofetilide(0) or DOFN) and cationic, i.e. positively charged (Dofetilide(+) or DOFC) dominant drug ionization forms (**Figure 2A**), which have different polarities, lipophilicities and lipid membrane permeabilities. This was demonstrated in our previous study on the example of another hERG blocking drug sotalol, which has similar functional groups and can also exist in cationic and neutral forms (34). Moreover, neutral and charged forms of a drug likely have different affinities for their protein targets, which was demonstrated in a recent study using hERG homology models and preliminary empirical force field models of dofetilide (33). Hence, for this study, we have developed and validated structural models for charged and neutral forms of dofetilide, compatible with biomolecular all-atom CHARMM force fields (see SI Methods). Our optimized DOFN and DOFC all-atom force field models are in good agreement with high-level quantum mechanical (QM) gas-phase molecular geometries, torsional energy profiles and interactions with water (see **SI Tables S1-S3, Figures S1-S2** and discussion therein).

**Figure 2.**
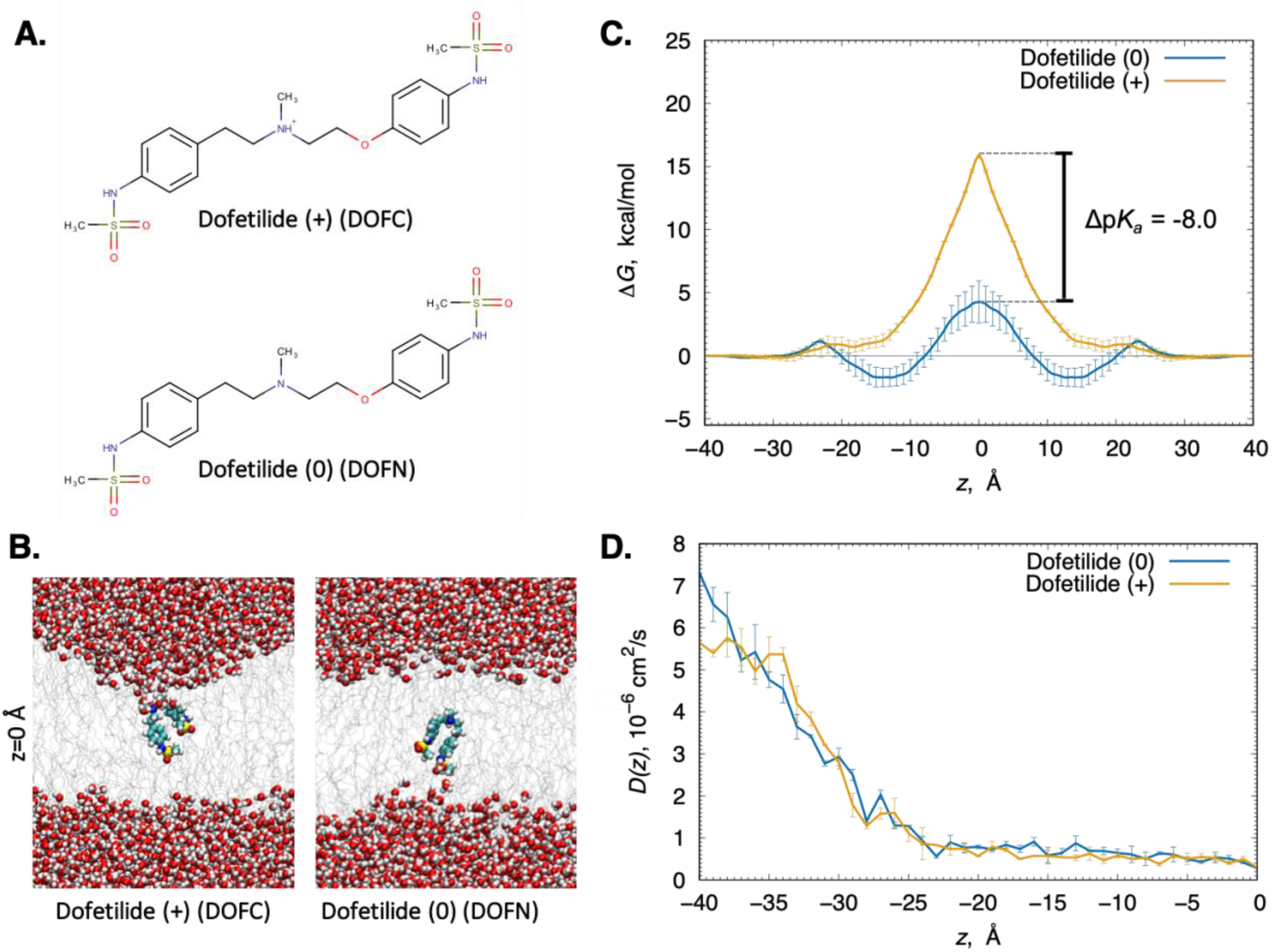
Translocation of charged and neutral dofetilide across a POPC lipid membrane. **(A)** Chemical structures of charged (top) and neutral (bottom) dofetilide. **(B)** Representative snapshots of drug in the membrane center (*z* = 0) from umbrella sampling simulations. Dofetilide and water molecule are in space-filling representation (C – cyan, O – red, N – blue, S – yellow, H – white), lipid tails are shown as gray sticks. **(C)** Potential of mean force (PMF) and **(D)** diffusion coefficient profiles of neutral (blue) and charged (orange) dofetilide crossing POPC membrane. Both PMF and diffusion coefficient profiles were symmetrized, and error bars were computed from profile asymmetries. p*K*_a_ shift at *z*=0 is shown by a black bar in panel **C** with the complete profile in **SI Fig. S4B**.

The derived parameters were validated by performing umbrella sampling MD simulations of DOFC and DOFN partitioning across a hydrated 1-palmitoyl-2-oleoyl-phosphatidylcholine (POPC) lipid bilayer. From these simulations, the drug water – membrane distribution coefficient log *D* (derived using Eq. 2), can be compared with experimental counterparts, as we have done previously (34). Moreover, drug–membrane interactions are crucial for understanding drug pharmacological properties, as cell membrane localization and permeation rates mediate the mechanism by which a drug affects its targets (34).

The results shown in **Figure 2B** demonstrate that there are substantial membrane perturbations as cationic dofetilide, DOFC, moves towards the membrane center, whereas this was not observed for the neutral species, DOFN. Therefore, it is not unexpected that the energetic barrier for DOFC crossing the membrane is high (Δ*G=* 15.8± 0.1 kcal/mol), and it is substantially lower (Δ*G=* 6.0± 1.8 kcal/mol) for DOFN (**Figure 2C**). Moreover, while the neutral form of the drug has interfacial binding wells of *–*1.7± 0.7 kcal/mol at |*z*|=14 Å, the charged one does not bind at the water-membrane interface at all (**Figure 2C**). A membrane crossing barrier almost 2.6 times higher for DOFC compared to DOFN indicates much faster rate of crossing for the latter, whereas interfacial binding of the neutral form signifies that it can accumulate in that region. Based on free energy differences of two forms, we computed how drug p*K*_a_ decreases as it moves across the membrane indicating its drastic deprotonation in the membrane interior (see **Figure 2C and SI Figure S4C**). The free energy profiles in **Figure 2C** and Eq. 2 were used to compute the water-membrane distribution coefficient, log*D* = 0.32 at pH 7.2, which agrees reasonably well with experimental values of 0.84, 0.98 derived from water-octanol (49, 50) and 2.08 from water -- Immobilized Artificial Membrane (IAM) (51) partitioning experiments.

Since we determined that DOFN accumulates at the water-membrane interface, we primarily expected it to be largely membrane-bound, contrary to its cationic counterpart DOFC. Indeed, our ∼2.5 μs long unbiased MD simulations demonstrate that the neutral form of the drug quickly (in <200ns) embeds into the membrane, while DOFC remains predominantly in aqueous solution (see **SI Figure S11A**). These results indicated that DOFN could potentially access hERG through a hydrophobic pathway, but that the charged form will only enter through the aqueous pathway (52). Moreover, due to a substantially smaller energetic barrier (**Figure 2C**), we expect much faster membrane translocation of DOFN. We also computed the diffusion coefficient, 𝒟(*z*), profiles across the POPC membrane for both forms of the drug, with values in the bulk water, 𝒟**_W_,** of ∼6×10^−6^ cm^2^/s for DOFC and ∼7×10^−6^ cm^2^/s for DOFN, each attenuating to 𝒟**_M_**<0.5×10^−6^ cm^2^/s at the membrane center (**Figure 2D** and **Table 1**). Such an order of magnitude drop in the diffusion in the membrane interior was observed in our previous studies for ions and other drugs as well (30, 34, 53). Based on these data, we estimated the membrane translocation rate of neutral dofetilide (see **SI Methods**) as 7.96 ± 1.37 ms^−1^. For a charged drug, its membrane crossing rate is expected to be 3-4 orders of magnitude smaller than that for the neutral one and thus is not expected to make a significant contribution to dofetilide membrane translocation.

### Open Conducting hERG Model

A crucial component of this work is a structural atomistic model of the hERG channel in a well-characterized, stable conformational state. Here we sought to develop an open, conducting model of the hERG based on the published cryo-EM structure (33). This was achieved by rebuilding missing loop residues using Rosetta (see **SI Methods**) thus constructing a complete model of the hERG pore domain (PD) and voltage sensing domains (VSD), and tested the ability of this channel model to conduct K^+^ ions in MD simulations. After an extended stage system equilibration (**see SI Table S4**), we performed a ∼1 μs-long unbiased MD simulation, which revealed that the PD remained open with a pore radius of ∼4 Å near activation gate, well hydrated and deviating from the initial structure by less than 3 Å (**SI Figure S5**). As expected, we did not observe ion conduction events in this simulation running at zero voltage.

However, we observed multiple K^+^ ion conduction events during an unrestrained, multi-μs MD simulation with an electric potential of 750 mV (positive inside) continuously applied along the *z* axis of the system, corresponding to the membrane normal, as shown in **Figure 3**. A representative molecular snapshot of this open conducting hERG model is shown in **Figure 3A**. The pore is open and solvent-accessible, while ions within the selectivity filter have adopted the canonical positions in an alternating “water-ion-water” orientation with ions at sites S1, S3 and cavity. Several frames from a representative conduction event of a K^+^ ion (brown ball) across the selectivity filter are shown in **Figure 3B**, with other ions also colored to match traces from **Figure 3C**. Ten conduction events through the channel were observed in ∼5 μs total simulation time (see **Figure S6A**), seven of which occurred within a 300 ns timespan, shown in **Figure 3C**. The ion SF transitions that facilitate conduction depicted by the brown ion are mediated by both water-mediated or “soft” (*t* = 478 ns) and direct or “hard” (*t* = 490 ns) knock-on events. K^+^ conduction ceases after 1.75 μs, coinciding with a marked reduction in the channel pore diameter and consequently pore dehydration (SI **Figures S6A and S7C).**

**Figure 3.**
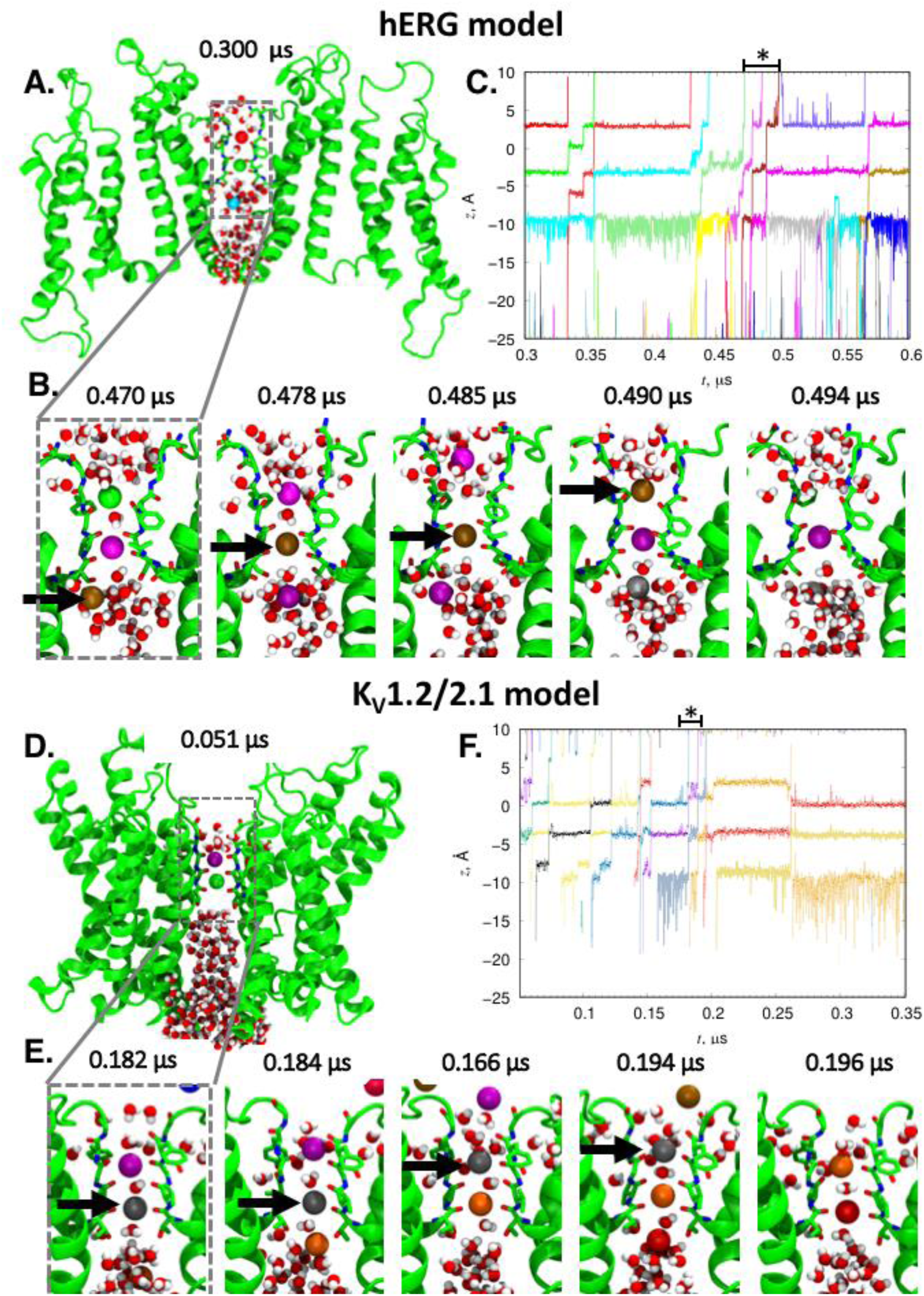
K^+^ conduction of open-state hERG and K_V_1.2/2.1 chimera channel models under an applied 750 mV voltage. 300 ns fragments of 5 μs trajectories, where most conduction events took place, are shown. See Figure 4 for complete trajectories. **(A, D)** Initial frames in these fragments (*t* = 0.300 and 0.051 μs, respectively) showing two opposite protein chains (green ribbons with SF S624-G628, S6 helix Y652 and F656 residues for hERG and SF T370-G374 residues for K_V_1.2/2.1 shown as sticks with red O and blue N), pore ions (colored spheres) and waters (red/white). **(B, E)** Close-up views of the channel SF in the same representation at different time points, showing a complete translocation of one ion (brown or gray ball, respectively), indicated by an arrow. **(C, F)** Time series of ion *z* positions (with respect to the SF backbone center of mass). Colors of the *z* profiles match those of the ions in panels **A**, **B** and **D**, **E**. Portions of the profiles corresponding to snapshots in panels **B** and **E** are indicated with an asterisk.

As a control, we also simulated the conduction of the K_V_1.2/2.1 paddle chimera (depicted in **Figure 3D**), well-studied previously (54, 55), under the same simulated conditions (750 mV applied voltage and 0.15 M KCl concentration). Our results indicated that the K_V_1.2/2.1 channel can also conduct multiple K^+^ ions during ∼5 μs long simulations (one such event is depicted in **Figure 3E**) and notably does not undergo hydrophobic pore collapse as in hERG (SI **Figures S6B and S8C).** We observed 9 conduction events within an initial ∼225 ns timespan (**Figure 3F)**; ∼70% more per unit time than in hERG conduction runs. Interestingly, this difference is in line with hERG and K_V_1.2 single-channel conductances observed experimentally (12.1-13.5 pS for hERG (56, 57) and 14-18 pS for K_V_1.2 (58)), although the small number of such events was not sufficient to draw any quantitative conclusions.

Moreover, we also observed long refractory periods in both hERG and the K_V_1.2/2.1 simulations (see SI **Figure S6**). They are correlated with pinching of the middle or top of the SF for hERG and K_V_1.2/2.1, respectively, which is evident by examining time series of distances between corresponding Gly residues (626 for hERG and 374 for K_V_1.2/2.1), depicted in **SI Figures S7B and S8B**. Interestingly, we observed top of the SF pinching and no ion conduction events in ∼2 μs long simulations of S641A mutant of hERG under 750 mV applied voltage (see SI **Figures S9B and S10C**). This mutant is known to facilitate channel inactivation (59), and thus such pinching could potentially be related to C-type inactivation, although simulation time scales are drastically smaller. Importantly, absence of conduction events for this hERG mutant indicates that even at high voltages used in our simulations we can distinguish between conducting and non-conducting channel states.

### Dofetilide Binding Pathways

Having developed dofetilide force field models and validated that our hERG open-state model remains open for at least 1 μs and can conduct K^+^ ions during that time, we sought to identify key drug-protein interaction pathways. To do this, we used so called drug “flooding” simulations, as in our previous studies (30, 35), where multi-μs-long unbiased MD runs were performed starting from a system with multiple dofetilide molecules placed in the aqueous solution around hERG, embedded in a POPC bilayer. For each system, we applied 25 mM aqueous concentration of the drug (corresponding to 20 dofetilide molecules) and ran simulations for 2.5 μs. The final systems are shown in **Figure 4A&B** for DOFN and DOFC, respectively.

**Figure 4.**
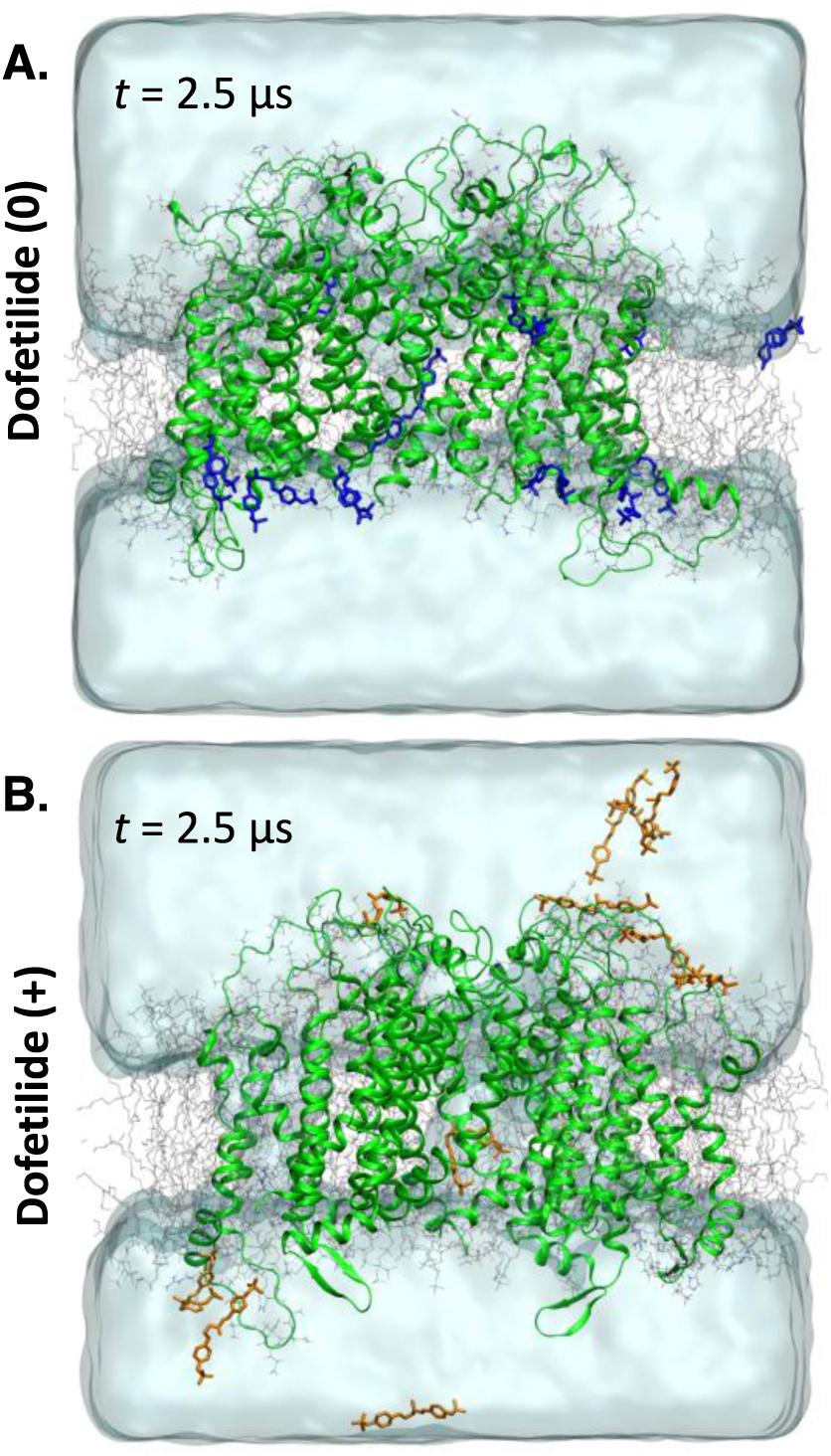
Unbiased hERG MD simulations in the presence of 25 mM initial aqueous dofetilide concentration. Snapshots of molecular systems at the end of 2.5 μs runs with neutral **(A)** and charged **(B)** dofetilide shown as blue and orange sticks, respectively. hERG is shown as green ribbons, lipid tails as gray sticks, water as aquamarine surface, K^+^ and Cl^−^ ions are not shown for clarity.

We observed that within first 500 ns of the simulation, most neutral drug becomes either embedded in the membrane or interacts with the protein (mostly at the intracellular side), thus reducing an effective aqueous concentration to less than 2 mM on average, while most DOFC molecules remain in aqueous solution transiently interacting with water-exposed hERG residues (**See Figure 4A&B and SI Figure S11A**). In these simulations we have not encountered DOFN or DOFC translocation into the channel pore via a lipophilic pathway, observed in our previous local anesthetic – Na_V_ channel simulations (30, 35). However, for both DOFN and DOFC, we observed one spontaneous drug diffusion into the channel pore through the open intracellular gate after ∼35 ns and ∼244 ns, respectively (**Figures 5A&B**). Once there, both forms remained bound for the duration of the simulation interacting, with S6 helix residues lining the pore, in particular, F656 and Y652, canonical hERG binding residues for dofetilide and most other hERG blocking drugs (60–62). In fact, mutation of these residues greatly reduces dofetilide hERG binding affinity, with Y652A mutations, for example, diminishing block by more than 50-fold (63) and F656V mutations reportedly diminishing dofetilide block by up to 100-fold (64). Interestingly, we observed different modes of interactions for DOFN and DOFC in the hERG pore (**Figure 5)**. DOFN adapted mostly extended conformation, stretching all the way from the bottom of the SF, where it interacts with S624, to the bottom of the S6 helix, interacting with S660. In contrast, DOFC adopts bent orientation and interacts mostly with canonical Y652, F656, and Q664 residues in the S6 helix. This does not exclude other possibilities since the simulation time might not have been long enough to sample other drug conformations. Such limited sampling might have been also enforced by a partial pore closure associated with some S6 helix bending during these simulations (**Figure 5A&B** and **SI Figure S11**).

**Figure 5.**
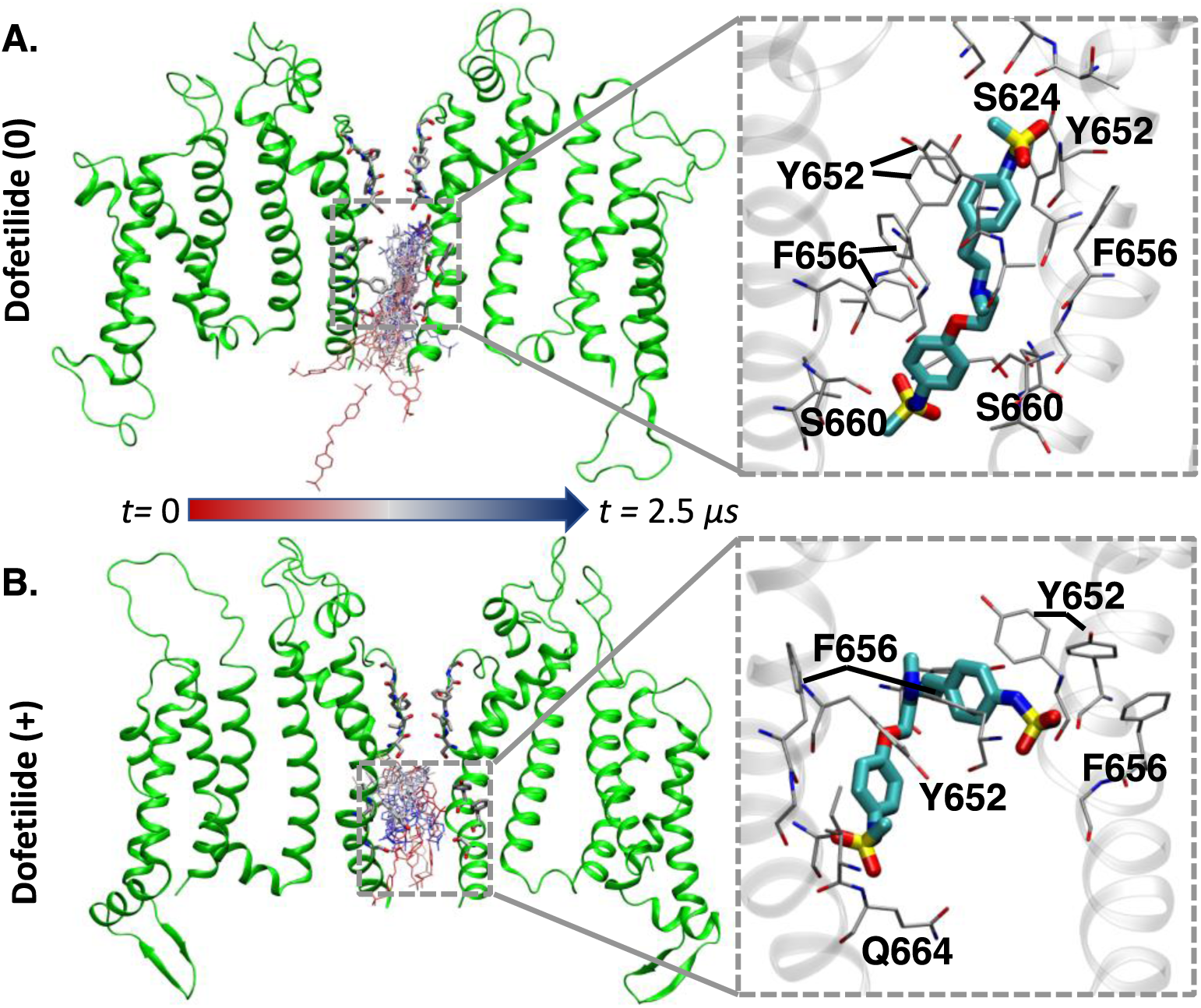
Dofetilide entry routes into hERG pore (left) and representative drug structures in the channel pore (right insets) from unbiased 25mM flooding simulations. *Left:* Snapshots of neutral **(A)** and charged **(B)** drug entering hERG pore from the beginning (red) to the end (blue) of 2.5 μs unbiased MD simulations. The final channel structure is also shown as green ribbons with SF S624-G628 and S6 helix Y652 and F656 residues shown as sticks with red O and blue N. *Right:* Insets demonstrate close-up view of representative neutral **(A)** and charged **(B)** drug structures in the channel pore. hERG backbone is shown as gray ribbons, interacting protein residues as thin sticks and dofetilide as thick sticks (C – cyan, O – red, N – blue, S – yellow).

### Energetics and Kinetics of Dofetilide Binding to hERG

Our drug flooding simulations demonstrated that both DOFC and DOFN likely access hERG pore through the aqueous intracellular gate. However, since we only observed singular drug ingress events and no instances of egress in each case, we were unable to quantitatively estimate energetics and kinetics of drug binding from these runs. Therefore, US MD simulations for drug – channel interactions were employed to compute these quantities along the channel pore.

A crucial prerequisite for US MD simulations is the generation of initial structures. Since dofetilide is a long and flexible molecule, sampling its re-orientation and conformational transitions in the narrow hERG channel pore is an extremely challenging task, as the results of US MD simulations can be greatly influenced by the initial drug orientation and conformation. To minimize this initial bias, and at the same time, make the initial system generation robust, we developed the strategy depicted in **Figure 6**. We employed steered molecular dynamics (SMD) to pull a drug molecule at a constant rate of 0.5 Å per ns from bulk aqueous solution (*z*=-50 Å) to the bottom of the SF (*z*=-5 Å) in five independent simulations, starting from different initial drug orientations. Since the optimal value for a pulling rate is known to be system specific (65) we chose one to allow for some drug relaxation within the channel pore. To seed our US MD simulations, we then randomly chose a frame from each of those 5 runs for the corresponding *z* position (**Figure 6B&C**), such that no two adjacent US windows were from the same pulling run. In all pulling and US MD runs we had weak harmonic restraints on the channel pore domain C_α_ and SF backbone atoms to eliminate channel transition to a different conformational state (See **SI Methods** for additional details).

**Figure 6.**
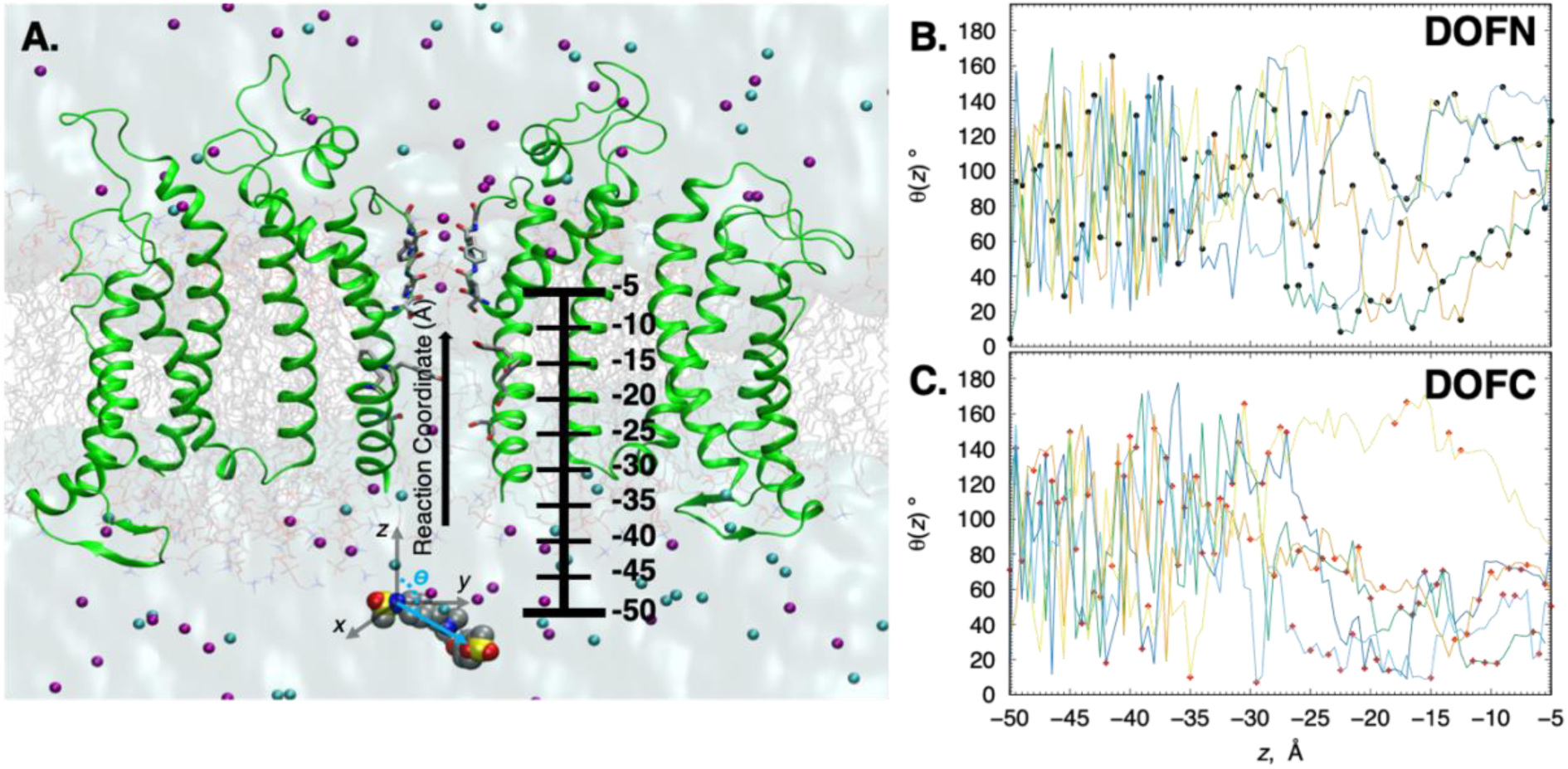
System setup for enhanced sampling MD simulations of dofetilide binding to hERG. *Left:* **(A)** Molecular snapshot of a simulation system with neutral dofetilide in bulk water moving into channel pore. hERG channel shown as green ribbons is embedded in hydrated lipid membrane with water depicted as aquamarine surface, POPC lipids as grey sticks, K^+^ and Cl^−^ ions as purple and cyan spheres. Neutral dofetilide is shown in the space-filling representation (C – gray, O – red, N – blue, S – yellow). Polar angle *θ* of dofetilide molecular vector, connecting two N atoms of sulfonamide groups, with the *z* axis is shown as well. The reaction coordinate for these simulations is the *z*-axis leading from the intracellular bulk aqueous solution (*z*=-50 Å) to the bottom of the selectivity filter (z=-5 Å) with the SF backbone center of mass used as a reference (*z*=0). *Right:* Neutral **(B)** and charged **(C)** dofetilide polar angle *θ* variation in drug pulling simulations is shown by colored lines, whereas *θ* values of structures from those simulations chosen to seed umbrella sampling runs are shown by black and red dots, respectively.

Potential of mean force (PMF) profiles computed from our US MD trajectories using the channel pore axis as the reaction coordinate allowed us to estimate the free energies of drug binding, Δ***G*_pore_**, and compute (using **Eq.3**) the dissociation constants (*K*_D*x*_) of dofetilide charged and neutral forms to the hERG channel in the open state. The results are summarized in **Table 1**, and in **Figure 7**, which shows that DOFC binds to the open state with Δ***G*_pore_** of about −5.9±0.4 kcal/mol at *z*=-20 Å, below the ring of S6 Y652 residues. This is in contrast with the neutral form of dofetilide, DOFN, which shows a much stronger binding with free energy of about −9.6±0.6 kcal/mol at *z*=-15 Å, above the Y652 ring and also interacting with S624 residues in the SF. DOFN binding is characterized by a fairly broad binding surface close to the SF region indicating that there are multiple binding poses possible, which is unlike the well-defined binding shown for charged dofetilide (**Figure 7A**). Importantly, for both DOFC and DOFN we observed that there are no free energy barriers for binding from bulk aqueous solution, which suggests that channel ingress kinetics (or “on” rate, *k*_on_) will be diffusion limited. In addition, since neutral dofetilide binds more favorably in the channel pore than its charged counterpart, we expect a strong downward shift in the drug p*K*_a_, around the binding site (*z* ≈ −20 to −15 Å) as was observed in the membrane interior (cf. **SI Figures S13B and S4B**).

**Figure 7.**
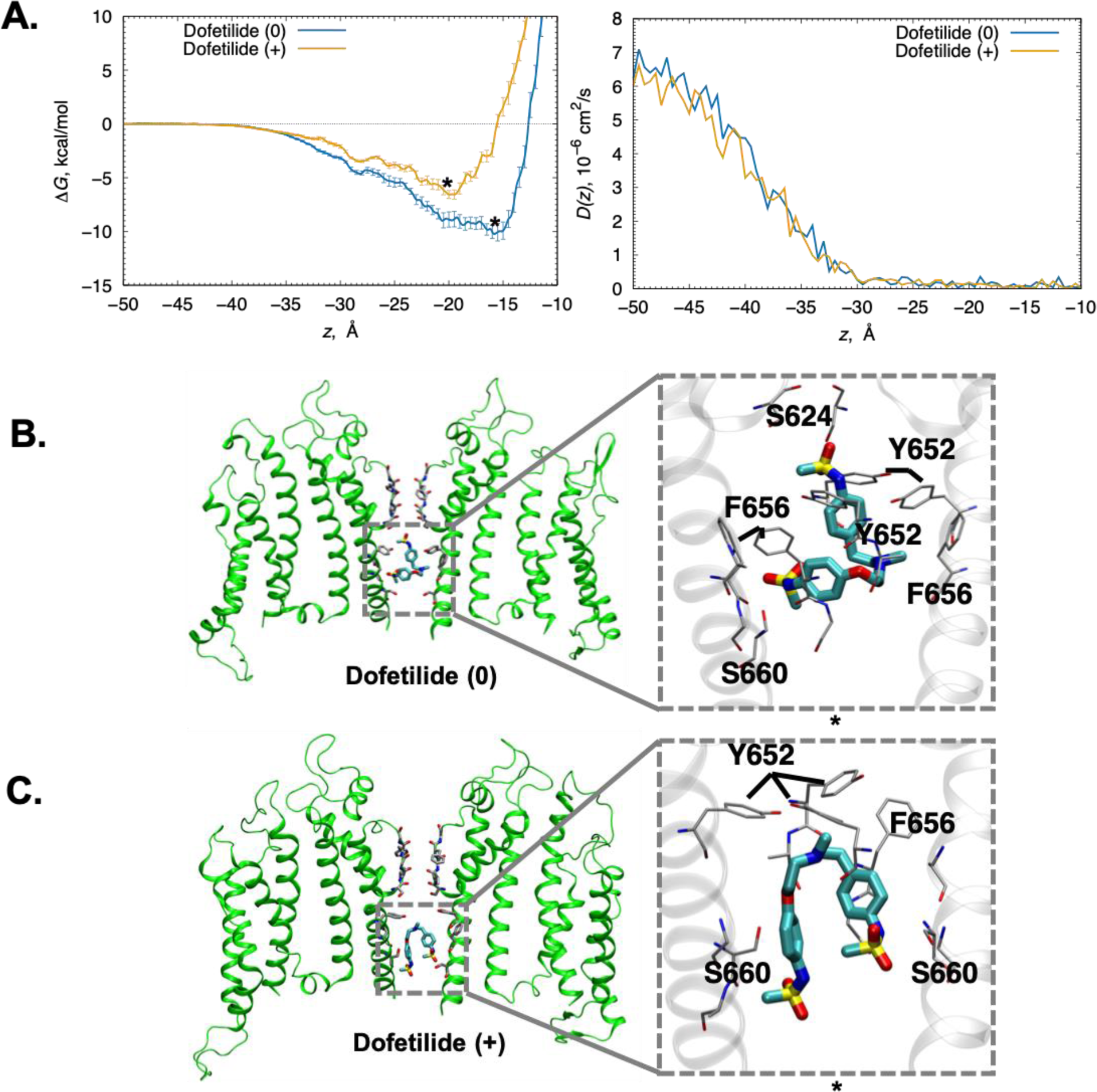
Charged and neutral dofetilide hERG binding from umbrella sampling MD simulations. **(A)** Free energy or PMF profiles (left) and corresponding diffusion coefficient profiles (right) for charged (orange) and neutral (blue) forms of dofetilide computed from umbrella sampling MD simulations. Error bars are computed from 5-ns block averaging and represent standard errors. Drug *z* positions of the representative lowest free energy snapshots in panels **B** and **C** are shown by asterisks. **(B, C)** Representative lowest free energy drug binding configurations for neutral **(B)** and charged **(C)** dofetilide in the hERG pore. Two opposite hERG chains with a drug bound are shown on the left, whereas insets on the right show close-up views of hERG residues interacting with dofetilide. hERG is shown as ribbons (green or gray), interacting residues as thin sticks and dofetilide as thick sticks (C– cyan, O – red, N – blue, S – yellow).

The equilibrium dissociation constant *K*_D*x*_ for DOFN and DOFC were computed to be 0.16 μM and 65.09 μM, respectively (**Table 1**). Taking into account the relative fractions of charged (38%) and neutral (62%) dominant drug ionization forms at pH 7.2, using its basic amine literature p*K*_a_ value of 7.0 (48), the overall *K*_D_ is estimated (using **Eq. 4**) to be ∼25.3 μM, which agrees favorably with IC_50_ values (3.5 μM, 10 μM, 11 μM) measured in electrophysiology experiments involving non-inactivating mutants of the channel or pulsing conditions disfavoring inactivated state (37, 57, 66). Other dofetilide p*K*_a_ values are available in the literature, such as e.g. 7.89±0.03 (67) and 8.0 (68), which would lead to overall *K*_D_ estimates of 54.1 or 56.2 μM, respectively, still in reasonable agreement with experiment.

The topologies of the binding pockets for the neutral and charged forms of dofetilide in representative poses are shown in **Figure 7B&C**, respectively. DOFN binds with one end pointing up toward polar residues in the SF, stabilized by interactions near the its base (S624) as well as a cluster of residues from S6 helix including S660, Y652 and F656 from multiple chains (**Figure 7B**), all residues that have been implicated in drug binding (62). One of the sulfonamide oxygens formed hydrogen bonds with S624, and the other sulfonamide group is hydrogen bonded to S660 (**Figure 7B**). These interacting residues are similar to ones sampled in drug “flooding” simulations discussed above. However, the drug conformation is different – extended in “flooding” simulations and bent in US MD runs (cf. **Figure 7B and Figure 5A**). One reason for this difference might be that in the former, the S6 pore helices were permitted to remain flexible, and adopt more constricted conformations. Whereas in US simulations we weakly restrained the S6 helix backbone to ensure a consistent conformational channel state, in which the pore cavity remains slightly more widened than in unbiased MD runs. In both cases, however, polar methanesulfonamide termini are stabilized by hydrogen bonding interactions with S624 and S660 residues. Interestingly, the S660 residue is known to be important for binding other bulky drugs like cisapride, terfenadine and ibutilide (69)

DOFC was observed to bind below the Y652 ring at z=-20 Å and is also coordinated by hydrophobic residues F656 and Y652 from multiple chains, with both terminal methanesulfonamide groups coordinated via hydrogen bonds with S660 residues (near the intracellular ends of S6 helices) from two chains (see **Figure 7C**). In other words, both ends of the molecule point down toward the bulk solvent, whereas its cationic ammonium group in the middle points up towards the SF and forms a hydrogen bond with Y652. Such drug conformation is fairly similar to one of those adopted by DOFC in drug “flooding” simulations (see **Figure 5**).

To investigate the stability of the drug binding poses observed in US MD simulations, we performed 1 μs long unbiased simulations, starting with the end frame of the US MD runs at *z*=-15 Å for DOFN and *z*=-20 Å for DOFC, in their putative binding sites (see **Figure 8**). DOFN remained bound in the pore and kept its bent conformation (**Figure 8A&C**). However, it tumbled with respect to the *z* axis (from ∼20 to 100° and back), and its *z* position fluctuated from −15 to −22 Å (**Figure 8A&B**), corresponding to a shallow trough at the US MD derived free energy profile (**Figure 7A**). An extended DOFN conformation, observed in our drug “flooding” MD runs, was not observed in these simulations which likely indicates existence of multiple stable binding poses with slow interconversion. Interestingly, for DOFC we observed several transient drug egress events for tens of ns, followed by its re-entry (**Figure 8A**). This was accompanied by drug tumbling with respect to the *z* axis (from ∼60 to ∼160° and back, see **Figure 8B**) and more importantly, its adoption of an extended conformation after ∼400 ns of the simulation (**Figure 8C&D**). This binding pose resembles one from DOFN “flooding” simulation (**Figure 5A)** but was observed in here in our DOFC stability test run. As for DOFN, this likely indicates existence of multiple low-energy binding poses separated by substantial barriers, precluding rapid interconversion.

**Figure 8.**
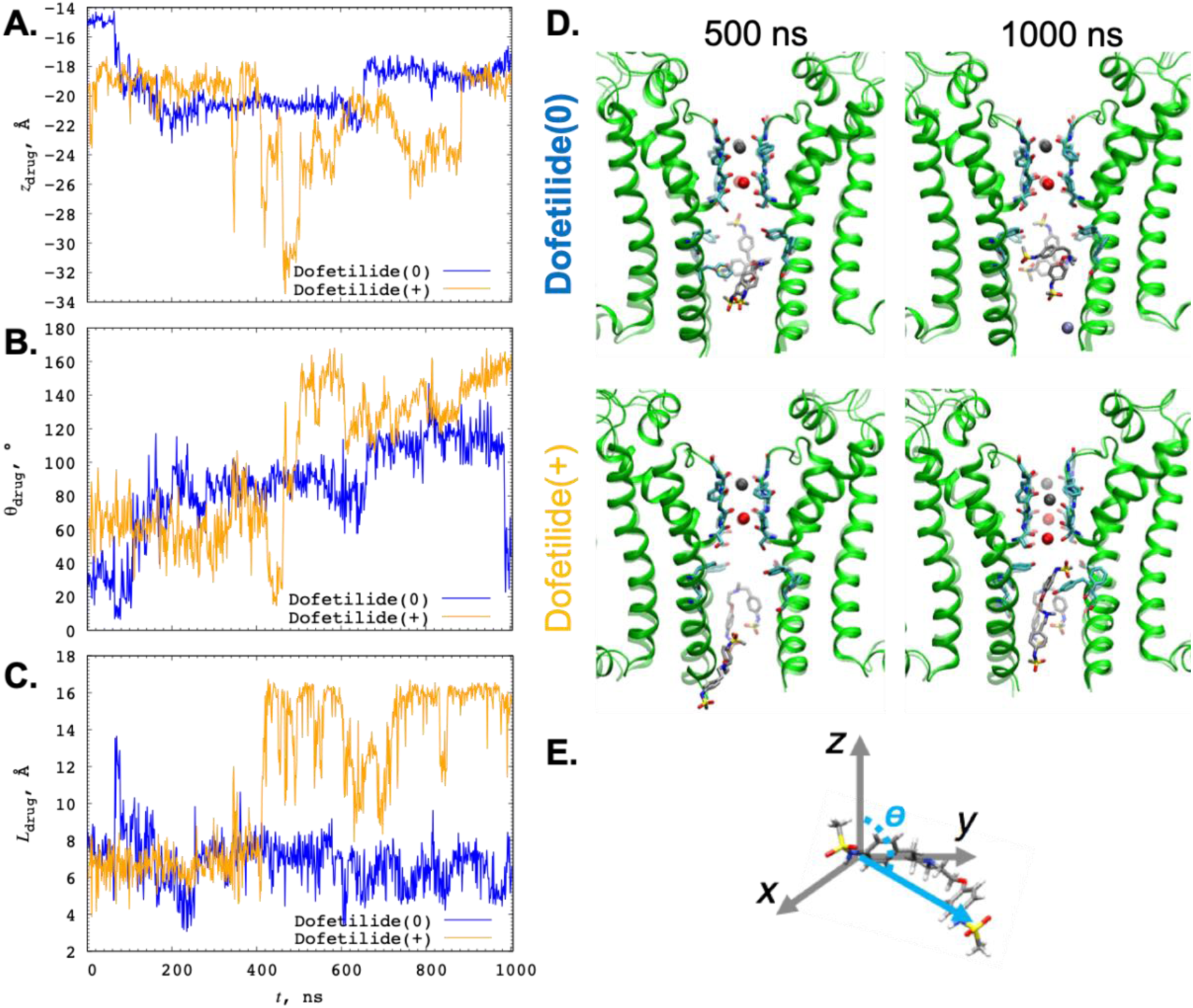
Stability of hERG drug-bound states identified from US MD simulations. **(A)** Time series of *z* position of dofetilide center of mass (COM) with respect to SF backbone COM beginning with the bound states (*z*=-15 Å for neutral and *z*=-20 Å for charged drug, respectively (see Figure 7A). **(B)** Drug tumbling with respect to *z*-axis, (polar angle *θ* between z-axis and vector connecting dofetilide N2 and N3 sulfonamide nitrogen atoms is illustrated in panel **E**), (**C)** Dofetilide molecule length measured as distance between N2 and N3 sulfonamide nitrogen atoms. (**D)** Molecular snapshots of dofetilide orientation in the hERG pore, with snapshots at 500 ns and 1000 ns shown in solid, and initial drug orientation from US MD simulations shown as semi-transparent.

We also computed the *z*-dependent diffusion profiles into the hERG pore for both drug forms (Figure 7A). Diffusion coefficients in the bulk aqueous solution, 𝒟**_W_,** are ∼6 ×10^−6^ cm^2^/s (at *z* = −50 Å) for both cationic and neutral dofetilide. They attenuate significantly to 𝒟**_pore_** of 0.11 or 0.15×10^−6^cm^2^/s for DOFN and DOFC, respectively (see **Table 1**), near the drug binding sites (i.e. at z ≈ −15 or −20 Å for DOFN and DOFC). Comparing the diffusion coefficient profiles for drugs in the channel pore and membrane interior (see **Figure 2D**), we see similar qualitative behavior – that their diffusion significantly slows down (by an order of magnitude or more) compared to bulk aqueous solution, due to a more viscous medium (in the membrane interior) or constricted space (in the channel pore). Interestingly, such reduction is more pronounced in the hERG pore (40-fold reduction) compared to the membrane interior (20-fold reduction).

Using the free energy (PMF) and diffusion coefficient profiles in **Figure 7A** along with **Eqs**. **5** and **6** we computed drug ingress rates (*k*_on_=666 μM^−1^s^−1^ for DOFN, and *k*_on_=534 μM^−1^s^−1^ for DOFC) and egress rates (*k*_off_=106 s^−1^ for DOFN and *k*_off_=3.47×10^4^ s^−1^ for DOFC) for open hERG pore, summarized in **Table 1**, which we will be using *directly* in our multi-scale computational model for predictive cardiac safety pharmacology, discussed in our companion paper (47). Since drug – channel association or ingress rates (*k*_on_) are diffusion-controlled, our estimates are similar for charged and neutral forms of the drug. The drug – channel dissociation or egress rates, *k*_off_, on the other hand, are directly proportional to drug dissociation constants, *K*_D*x*_. Therefore, stronger DOFN binding leads to a ∼65-fold reduction in its egress rate compared to DOFC. This also correlates with the results of our 1 μs long unbiased MD simulations of drug binding, which demonstrated partial drug unbinding in DOFC but not DOFN run.

We can also compare drug ingress rates at maximum physiological dofetilide plasma concentration *C*_max_ of ∼6.16 nM (see the companion paper (47)), *x· C*_max_ *· k*_on_ (where *x* is DOFN or DOFC fraction at pH = 7.2), equal to 2.5 and 1.2 s^−1^ for DOFC and DOFN, respectively, with that for DOFN membrane translocation, 7.96 ms^−1^ or 7,960 s^−1^. The latter is over 3 orders of magnitude faster indicating that kinetics associated with open hERG – dofetilide interaction will be a rate-determining step and should definitely be taken into account in our functional models, discussed in our companion paper (47).

## Discussion

In this study we take the initial steps towards a multi-scale modeling framework to predict the effect of dofetilide on the heart starting from atomic scale structural models of hERG – drug interactions. State-specific drug – channel ingress and egress rates computed from the atomic scale MD simulations were used to parameterize a functional kinetic channel model that captures the dynamical interactions of dofetilide with hERG (see **Figure 1B**) in the second part of this study (see the companion paper (47)). Our functional kinetic protein-scale model has been integrated into predictive functional kinetic models at the cell and tissue scales to expose fundamental arrhythmia vulnerability mechanisms and complex interactions underlying emergent behaviors.

Typically, experimental biophysical and biochemical data are used to predict parameters associated with drug effects on protein targets for functional multi-scale models (70). However, many important variables such as a particular drug ionization and channel conformation are difficult to control experimentally (71, 72). Atomistic MD simulations are ideally suited to address this challenging problem by providing necessary spatial and temporal resolutions to compute state-dependent drug binding affinities and rates.

In this study we developed and validated models of charged (cationic) and neutral forms of dofetilide, compatible with biomolecular CHARMM force fields, that were optimized using high-level QM data as a reference. Then we performed membrane partitioning MD simulations, revealing that only the neutral form of the drug is expected to localize at water-membrane interface, with a relatively fast translocation rate of ∼8 ms^−1^. The charged form is not expected to contribute to drug membrane translocation due to ∼4 times larger barrier. Our computed water – membrane distribution coefficient log*D*_MW_ of 0.32 at pH 7.2 is in reasonable agreement with experimental log*D* values for water-octanol of 0.84 – 0.98 (49, 50) and for water – artificial supported membrane of 2.08 (49), somewhat underestimated compared to the latter. The discrepancy does not necessarily indicate model deficiencies, and can be related to experimental methodology or data interpretation (73). There is also uncertainty in dofetilide p*K*_a_ value (48, 67, 68), which would affect computed log*D*_MW_ as well as composite drug-channel composite *K*_d_ value (see below), but not individual estimates for charged and neutral drug forms, used in our multi-scale functional model.

We also developed and validated a model of hERG in the open conducting conformation based on the recently published cryo-EM structure of hERG (33). The model demonstrated multiple K^+^ conduction events in ∼0.3 μs span of simulations under a strong +750 mV voltage, which was previously used to facilitate ion conduction for KcsA and K_V_1.2/2.1 channels (54, 74). Qualitatively similar conduction behavior was observed in our reference K_V_1.2/2.1 channel simulation at the same simulation conditions. Importantly, we demonstrated that our hERG model likely represents an open conducting state, and that its alteration, via inactivation-accelerating S641A mutations, can render it non-conductive.

Having developed and validated atomistic models of dofetilide and hERG, we started studying their interactions via all-atom MD runs. First, we performed so called multi-microsecond drug “flooding” MD simulations with multiple drug molecules, which indicated that both charged and neutral dofetilide can access the channel pore through the intracellular gate and form long-lasting interactions with canonical drug binding residues (F656 and Y652). Notably, none of the dofetilide molecules moved into the pore via a lipophilic pathway, in agreement with recent lipid-facing hERG mutant screening simulations (52), but observed for local anesthetic drugs binding to Na_V_ channels (30, 35). These simulations provided interesting qualitative insights and indicated the need to use enhanced sampling techniques to obtain quantitative state-specific drug binding energetics and kinetics.

To obtain dofetilide free energy and diffusion coefficient profiles, we performed umbrella sampling MD simulations of charged and neutral dofetilide binding to hERG pore via the intracellular gate (see **Figure 7A**). We observed stronger binding of neutral dofetilide deep inside the channel pore and close to the bottom of the SF compared to charged dofetilide, which was bound less tightly further down the channel pore. This suggests possibility of drug deprotonation inside the pore, although not as drastic as in the membrane interior. Interestingly, we observed that drug diffusion is slowed down by an order of magnitude in both channel pore and membrane interior, but more though in the former due to constricted environment (**Figures 7A and 2D**). Based on the free energy and diffusion data we computed drug dissociation constants as well as drug ingress (entry) and egress (exit) rates (see **Table 1**), which were incorporated into our multi-scale functional model, as discussed in the companion paper (47). As expected, the diffusion-controlled drug ingress rates (*k*_on_) are similar for both charged and neutral dofetilide species, whereas egress rate (*k*_off_), dependent on drug binding affinity, is almost 2 orders of magnitude lower for neutral dofetilide. Importantly, dofetilide-hERG ingress rate values at maximum physiological plasma concentration are over 3 orders of magnitude smaller than our estimate for neutral dofetilide membrane permeation rate, indicating that drug-channel association is likely to be a rate-limiting step for the entire drug/channel/membrane system and should be incorporated into our functional kinetic channel model (see **Figure 1B** and companion paper (47)). Our computed composite *K*_d_ value estimates of 25 - 56 μM, taking into account neutral and charged dofetilide contributions at pH 7.2 and p*K*_a_ of 7.0 - 8.0 (48, 67, 68), are in reasonably good agreement with experimental IC_50_ estimates (3.5 - 11 μM) of dofetilide binding to non-inactivated channel state (37, 57, 66). This indicates that our atomistic channel – drug model is likely to be accurate enough to be used in providing parameters for our predictive multi-scale kinetic models of cardiac safety pharmacology.

Notably, experimental data suggest that dofetilide binds considerably stronger to inactivated hERG state, with IC_50_ in the low nanomolar range (36, 75-78). In this study we considered drug binding only to an open conducting channel state, for which we have structural data. In order to include dofetilide binding to the inactivated state into our functional model for this prototype study, we used the experimental ratio of binding affinities to these two states (see the companion paper (47)). However, future model development will necessitate a structural model of the channel inactivated state (see SI for a more detailed description).

The atomistic molecular modeling and simulation protocol we have developed in this work was utilized for estimating multi-scale functional model parameters as described in the companion paper (47). It can be used, for example, as a structural predictor of experimental electrophysiology channel inhibition or drug radioligand binding assays. It is modulatory and can be easily modified or amended as needed, and steps can be potentially automated using computational workflows, as was done recently for our functional models (79). We are working to expand these studies to other hERG blocking drugs with different pro-arrhythmia proclivities and also to consider multi-channel block as well, as emphasized in CiPA initiative (80, 81). This study represents a crucial step towards our ultimate goal to provide and thoroughly test a transferrable *in silico* protocol for accurate and efficient prediction of the cardiotoxicity of any drug or drug candidate based on their chemical structures, which will help decrease costs associated with drug development and lead to development of safer drugs and save human lives.

### Materials and Methods

#### Drug Force Field Parameterization

Atomistic model parameters for charged (DOFC) and neutral dofetilide (DOFN) were based on generalized CHARMM force field (CGENFF) program (82, 83) initial guesses and then optimized using a standard set of quantum mechanical (QM) calculations utilizing the ffTK plugin (84) for the Visual Molecular Dynamics program (VMD) (85).

#### Ion Channel Models

A recently published cryo-EM structure (PDB ID: 5VA2) (23) truncated at residues <405 and >668 was used a template for an open state hERG channel model, thus comprising the voltage sensing domain (VSD) and pore domain (PD) of the channel. Rosetta structural modeling software with symmetry-imposed local iterative refinement and fragment-based loop modeling protocols (86–88) was used to generate the missing loop regions. For an open state K_V_1.2/2.1 model, VSD and PD of the X-ray structure (PDB ID: 2R9R) (89) were used. See **SI Methods** for more details.

#### MD Simulation Setup

CHARMM-GUI toolkit (90) and NAMD (91) program were used in order to build and equilibrate the simulation systems, which included 1-palmitoyl-2-oleoylphosphatidylcholine (POPC) lipid bilayer with or without embedded ion channel protein, oriented along the *z* axis and solvated by 0.15M aqueous KCl solution. Hydrated POPC bilayers were built around dofetilide molecules placed at different *z* positions in drug-membrane translocation runs. In simulations containing ion channel and drug molecule(s), the latter were initially placed in bulk aqueous solution. All the simulations were run at 310 K and 1 atm pressure (when applicable).

#### Ion conduction and drug flooding channel simulations

hERG systems were equilibrated with NAMD for 90 ns using gradually reduced restraint regime (see SI Methods and SI Table S4) and then simulated on the Anton 2 supercomputer for 2.5 – 5 μs. K_V_1.2/2.1 systems were equilibrated with NAMD for 50 ns before 5 μs Anton 2 production runs. Ion conduction simulations contained no drug and had +750mV transmembrane voltage via applied electric field across the simulation box *z* axis and were run using the *NVT* ensemble. Drug flooding simulations were set up such that there was a 25 mM initial concentration of dofetilide randomly distributed in aqueous solution and were run in the *NPT* ensemble.

#### Umbrella sampling (US) MD simulations

US MD simulations (46) were used to sample drug translocation across a POPC membrane and into the hERG channel pore, and potential of mean force (PMF) profiles were computed using weighted histogram analysis method (WHAM) (92). Error bars were estimated as standard errors of mean from block averaging of PMF data. For drug-membrane partitioning simulations, there were 81 independent US windows with 2.5 kcal/mol/Å^2^ harmonic restraint in 1 Å intervals along membrane normal (*z* axis) for −40 Å ≤ *z* ≤40 Å with respect to center of mass (COM) of the membrane. For hERG-drug US MD simulations there were 91 independent US windows with 10.0 kcal/mol/Å^2^ harmonic restraint, spaced in 0.5 Å intervals for −50 Å ≤ *z* ≤ −5 Å with respect to hERG selectivity filter (SF) C_α_ atoms. The pore domain C_α_ and all SF backbone non-H atoms were subject to 0.2 kcal/mol/Å^2^ harmonic restraints. Cylindrical flat-bottomed restraint with 10 Å radius was used to restrict sampling to the channel pore. All US MD simulations were run in NAMD.

#### Drug – membrane partitioning thermodynamics and kinetics

Dofetilide – POPC membrane partitioning coefficients *P_x_* for charged (*x*=1) and neutral (*x*=0) drug forms were computed from corresponding US MD simulations as done previously (34):

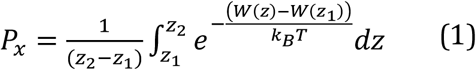

where *z*_1_ *and z*_2_ represent points in the bulk aqueous solution on opposite sides of the membrane bilayer, *k*_B_ is the Boltzmann constant, *T* is the temperature in Kelvin, and *W*(*z*) is the PMF along the drug – membrane reaction coordinate. The distribution coefficient, log *D*, was computed as a composite value:

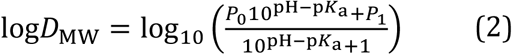

where p*K*_a_ is acid dissociation constant of dofetilide. Drug membrane translocation rate was estimated using Kramer’s rate equation, as was done previously (30).

#### Drug – channel binding thermodynamics and kinetics

The effective equilibrium dissociation constants *K_x_* of charged (*x*=1) and neutral (*x*=0) drug forms were computed from PMFs of dofetilide – hERG pore binding US MD simulations as done previously (33):

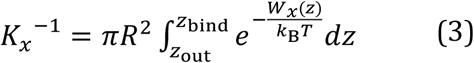

where *R* is the restraint radius of the drug that encompasses the width of the channel pore (10Å), *k*_B_ is the Boltzmann constant, *T* is the temperature in Kelvin, and *W_x_*(*z*) is the potential of mean force (PMF) along the reaction coordinate from a point in the intracellular bulk aqueous solution (*z*_out_) to a point below channel SF (*z*_bind_). The composite *K*_D_ value to be compared with available experimental data (see Table 1) was computed as:

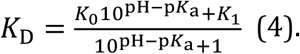

**Table 1.** Dofetilide membrane partitioning and hERG binding energetics.

|  | MD Simulations |  | Experiment |
| --- | --- | --- | --- |
|  | DOFN | DOFC |  |
| **Lipid membrane partitioning data** |  |  |  |
| $\log P_x$ | $0.49 \pm 0.14$ | $-0.32 \pm 0.08$ | — |
| $\log D_{MW}$ or $\log D_{ow}$ | $0.32 \pm 0.13$ | | $0.84^a, 0.96^b, 2.08^c$ |
| $D_w$ ($10^{-6} \text{ cm}^2 \text{ s}^{-1}$) | $6.34 \pm 0.97$ | $5.59 \pm 0.66$ | — |
| $D_M$ ($10^{-6} \text{ cm}^2 \text{ s}^{-1}$) | $0.44 \pm 0.15$ | $0.43 \pm 0.09$ | — |
| **hERG pore binding data** |  |  |  |
| $D_{\text{pore}}$ ($10^{-6} \text{ cm}^2 \text{ s}^{-1}$) | 0.11 | 0.15 | — |
| $\Delta G_{\text{pore}}$ (kcal mol$^{-1}$) | $-9.64 \pm 0.63$ | $-5.94 \pm 0.43$ | — |
| $K_x$ ($\mu\text{M}$) | $0.16 \pm 0.09$ | $65.1 \pm 31.3$ | — |
| $K_D$ or $IC_{50}$ ($\mu\text{M}$) | $25.3 \pm 12.1$ | | $3.5^d, 10^e, 11^f$ |
| $k_{\text{on}}$ ($\mu\text{M}^{-1}\text{s}^{-1}$) | 666 | 534 | — |
| $k_{\text{off}}$ ($\text{s}^{-1}$) | 106 | 34700 | — |

Water-membrane partitioning coefficients ( $\log P_x$ ) and distribution coefficient ( $\log D_{MW}$ ) at pH=7.2 as well as diffusion coefficients in bulk water ( $D_w$ ) and membrane center ( $D_M$ ) were computed from 1D umbrella sampling MD simulations for charged (DOFC) and neutral (DOFN) dofetilide translocation across a POPC membrane. They were compared to experimental water-octanol distribution coefficients ( $\log D_{ow}$ ) from references <sup>a</sup> (50) and <sup>b</sup> (49) and water – immobilized artificial membrane (IAM) high-performance liquid chromatography (HPLC) distribution coefficient ( $\log D_{MW}$ ) from reference (50). Free energies ( $\Delta G_{\text{pore}}$ ) and diffusion coefficients ( $D_{\text{pore}}$ ) of DOFC and DOFN at the binding site in the channel pore as well as corresponding dissociation constants ( $K_x$ ), association ( $k_{\text{on}}$ ) and dissociation ( $k_{\text{off}}$ ) rates were computed from 1D umbrella sampling simulations along the channel pore. Net (composite) dofetilide dissociation constant ( $K_D$ ) at pH=7.2 was compared to the experimental half maximal inhibitory concentrations ( $IC_{50}$ ) of the drug to non-inactivated hERG from references <sup>d</sup> (66), <sup>e</sup> (57), <sup>f</sup> (37). See text for more details.

The rate constant for drug – channel binding, *k*_on_, was computed using a formulation of the Debye-Smoluchowski equation (93, 94):

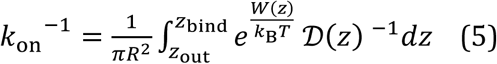

where *R* designates the radius of a cylinder (10 Å) that encompasses the channel pore, and 𝒟*(z*) is the local diffusion along the reaction coordinate. Having already computed *K*_D_ and *k*_on_, we can trivially compute *k*_off_ as:

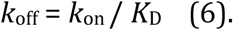

## ACKNOWLEDGMENTS

We thank Drs. Heike Wulff, Jon T. Sack, Kazuharu Furutani (UC Davis), Toby W. Allen, Celine Boiteux, Emelie Flood (RMIT University) and members of the C.E.C., V.Y.-Y., and S.Y.N. laboratories for helpful discussions. This research was supported by National Heart, Lung, and Blood Institute Grants U01HL126273, R01HL128537, R01HL128170, and OT2OD026580 (to C.E.C.), American Heart Association Predoctoral Fellowship 16PRE27260295 (to K.R.D.), NIGMS-funded Pharmacology Training program T32GM099608 (to J.R.D.D), American Heart Association Career Development Grant 19CDA34770101 (to I.V.) and UC Davis PMB Research Partnership Fund (to I.V. and C.E.C.). Anton 2 computer time was provided by the Pittsburgh Supercomputing Center (PSC) through Grant R01GM116961 from the National Institutes of Health and was awarded through PSCA17085P (to I.V. and C.E.C.), PSCA16108P (to C.E.C., I.V. and K.R.D.) allocations. The Anton 2 machine at PSC was generously made available by D.E. Shaw Research. Other computer time allocation was used through XSEDE award MCB170095 (to C.E.C., I.V. and K.R.D.), National Center for Supercomputing Applications (NCSA) Blue Waters Broadening Participation allocation (to C.E.C.), Compute Canada Research Allocation award (to S.Y.N.).

## FOOTNOTES

### Author contributions

K.R.D., V.Y.-Y.,S.Y.N., I.V., and C.E.C. designed research; K.R.D, J.R.D.D., P.-C.Y., S.B., V.A.N., V.Y.-Y., S.Y.N., C.E.C. and I.V. performed research and analyzed data; and K.R.D, J.R.D.D.,V.Y.-Y., S.Y.N., C.E.C. and I.V. wrote the paper.

