## Supporting Information for "Atomistic modeling towards predictive cardiotoxicity"

**AFFILIATIONS:**

[*] Correspondence:

### Supporting Information

#### SI Methods

##### General Methodology Overview

Atomistic molecular dynamics (MD) simulations allow for quantitative prediction for drug binding affinities, which can be checked against experimental data, as well as ligand – protein association and dissociation rates, which will be used in our multi-scale functional models of cardiac safety pharmacology, described in companion paper, Part 2. The special-purpose Anton 2 supercomputer from DE Shaw research (4), used in our study, allows for several microseconds of all-atom MD simulations to be run in less than a day, and GPU and CPU supercomputing clusters allow for enhanced sampling MD simulation methods (5, 6), such as umbrella sampling (US) (7, 8) used in our study, to be run efficiently in parallel. The general CHARMM force field for drug-like molecules (9) and corresponding program for atom type matching and parameter prediction (10, 11) enables the robust development of atomistic drug models that are compatible with protein, lipid membrane, water and ion CHARMM force field parameters (12). Umbrella sampling MD simulations of drug interactions with the lipid membrane allows validation of models through comparison between experimental and computed partitioning data (13). Multi-microsecond-long all-atom simulations under applied voltage permits the critical evaluation of ion conduction and channel model stability, and for the prediction of channel conformational states (14-17). Enhanced sampling MD simulations of protein – drug interactions can be used to determine free energy and position-dependent diffusion coefficient profiles, as was done previously (1-3), to compute binding affinities and drug ingress and egress rates.

##### Drug force field parameterization

The molecular structure of dofetilide was obtained from the ZINC database (accession number 49583080) (18). Initial parameters for charged (DOFC) and neutral dofetilide (DOFN) were based on general CHARMM (Chemistry at Harvard Molecular Mechanics) force field (CGENFF) initial guesses (10, 11) and parameters with poor chemical analogy were optimized following the suggested CGENFF force field methodology (9) using a standard set of quantum mechanical (QM) calculations utilizing the ffTk plugin (19) for the Visual Molecular Dynamics program (VMD) (20). Gas-phase quantum mechanical (QM) calculations utilizing Møller–Plesset (MP2) and Hartree-Fock (HF) perturbation theory and the 6-31(d) basis set in Gaussian 09 (21) program were used to compute target data for parameter optimization.

MP2/6-31G(d) molecular dipole magnitude and orientation as well as scaled HF/6-31G(d) interaction energies with water were used for the optimization of partial atomic charges compatible with the all-atom CHARMM biomolecular force fields (22). Internal bond and angle parameters were validated by comparison to MP2/6-31G(d) optimized geometries and scaled vibrational frequencies, and differences within 0.01 Å and 1° between QM and force field, i.e. molecular mechanical (MM) equilibrium bond and angle values were sought. Finally, the dihedral angle parameters were optimized to reproduce MP2/6-31G(d) potential energy scans for rotation around a particular bond.

Optimized charges (**Table S1**) provide a good agreement with QM target dipole values. The optimized MM dipole moments are larger in magnitude compared to QM MP2/6-31G(d) dipole moments by 19% (9.7 vs. 8.1 D) for neutral dofetilide and 17% (10.8 vs. 9.2 D) for charged dofetilide, which are close to the 20% lower-end threshold suggested for CGENFF The water interaction distances were all within 0.4 Å of QM target values (see **Tables S2 and S3**). Water interaction energies were also in good agreement with QM values, with root mean squared errors (RMSE) of 0.53 kcal/mol for neutral dofetilide, and 1.44 kcal/mol for charged dofetilide, respectively (**Table S2 and S3**). For neutral dofetilide, there were high penalty scores for bond angles involving the central amine group N atom, C1-N1-C11, C1-N1-C2, and C11-N1-C2, and optimization yielded an absolute difference of 0.35°, 0.02°, and 0.42°between MM and QM values, respectively. There were no high penalties for internal bond and angle parameters for charged dofetilide, thus the CGENFF program generated values were used. For neutral dofetilide, there were four high-penalty dihedral angle parameters, and for charged dofetilide there was one such parameter, which needed to be optimized. Dihedral angle parameter optimizations resulted in substantial improvement over CGENFF initial guesses, with optimized torsional energy minima within ~1 kcal/mol of QM values (see **Figure S2**). Final topology and parameters for neutral and charged dofetilide are provided at the end of this Supplement in SI Appendix.

**General MD Simulation Setup**

The CHARMM-GUI online toolkit (23), CHARMM (24, 25), NAMD (5), and Anton 2 software programs were used in order to build and simulate the molecular systems in this study. All of them contained 1-palmitoyl-2-oleoylphosphatidylcholine (POPC) lipid bilayer hydrated by a 0.15 M aqueous KCl solution. The membrane normal axis was aligned along the z-axis in all cases. The hERG channel was placed in the bilayer center with its aqueous pore aligned with the membrane normal. Drug flooding simulations were initialized with a 0.025 M aqueous concentration of dofetilide. All NAMD simulations apart from ion conduction simulations under applied voltage, were carried out in an *NPT* ensemble with 1 atm pressure maintained by Langevin piston barostat (26), and 310K, controlled by Nosé-Hoover thermostat (27, 28). MD simulations with an applied voltage to study ion conduction were carried out in the *NVT* ensemble. Tetragonal cells with periodic boundary conditions (PBC) were used in all the simulations, and the SHAKE algorithm (29) was employed to fix the bonds to all hydrogen atoms, allowing for the use of a 2 fs time step. Electrostatic interactions were computed via Particle Mesh Ewald (30), with a mesh grid of 1 Å.

**Membrane permeation simulations of the hERG blocker dofetilide**

*Setup for dofetilide-membrane US MD simulations:* In order to validate our drug models against available experimental data, we ran water-membrane partitioning simulations of both charged and neutral states of dofetilide, DOFC and DOFN, utilizing the umbrella sampling (US) methodology (7). Initial system setup scripts were generated with the CHARMM-GUI web toolkit (23) and were modified to build the hydrated drug-membrane systems, which consisted of 128 POPC lipids, approximately 7000 water molecules, 21 or 22 $K^+$ and 22 $Cl^-$ ions to ensure 0.15 M electrolyte concentration and overall electrical neutrality, and one drug molecule, totaling approximately 38,400 atoms. CHARMM36 lipid force field (31), TIP3P water model (32), standard CHARMM ion parameters (33) and CGENFF (9) compatible drug parameters developed in this work were used throughout all simulations.

These calculations were run using the NAMD 2.12 program (34), CHARMM36 all-atom force field and TIP3P water model (31, 33), at a constant pressure of 1 atm and at a physiological temperature of 310 K.

*Details of dofetilide-membrane US MD simulations:* For each dofetilide model (charged and neutral), 81 independent simulation windows were created in which the center of mass (COM) of a randomly oriented drug molecule was placed at 1 Å intervals from $z$= -40 Å to $z$ = 40 Å with respect to COM of the membrane. In addition, for the membrane-spanning central windows, $z$= -20 Å to $z$= 20 Å, additional simulations were performed, in which the drug was flipped about its $x$-axis in order to enhance sampling, and hence reduce asymmetries in the computed potential of mean force (PMF) profiles (see **Figure S3**). For neutral dofetilide, sampling in the membrane interior remained poor (see **Figure S3B**), so additional simulations of central membrane-spanning windows beginning with a uniform drug orientation were run for 20 ns each. For all US MD simulations, the COM of the drug was restrained along the $z$ axis with a force constant of 2.5 kcal/mol/Å$^2$, and an additional 5 kcal/mol/Å$^2$ cylindrical restraint was applied in order to prevent its drift in the $xy$ plane. Umbrella sampling simulations for charged dofetilide were run for 20 ns for each window, and the first 5 ns were discarded to account for equilibration. Neutral dofetilide simulations were run for 30 ns for randomly oriented and flipped windows, and for 20 ns for uniformly distributed windows, and for each US window information from the first 5 ns was again discarded to account for equilibration. Each NAMD US simulation of charged and neutral dofetilide was carried out in the *NPT* ensemble with 1 atm pressure maintained by Langevin piston barostat (26), and 310K, controlled by Nosé-Hoover thermostat (27, 28). Tetragonal cells with periodic boundary conditions (PBC) were used in all the simulations, and the SHAKE algorithm (29) was employed to fix the bonds to all hydrogen atoms, allowing for the use of a 2 fs time step. Electrostatic interactions were computed via Particle Mesh Ewald (PME) (30), with a mesh grid of 1 Å. The PMF profiles were computed using the weighted histogram analysis method (WHAM) (35) with error bars computed from asymmetries. Symmetrized diffusion coefficient profiles (-40 Å to 0 Å shown in main text **Figure 2D**) were obtained using Laplace transform of drug position autocorrelation function (36) as was described in our recent studies (37, 38), with error bars computed from asymmetries.

*Membrane Partitioning Coefficients:* Membrane partitioning coefficients $P_x$ were computed as was done previously (38, 39):

$$P = \frac{1}{(z_2 - z_1)} \int_{z_1}^{z_2} e^{-\frac{\{W(z) - W(z_1)\}}{k_B T}} dz \quad (S1)$$

where $W(z)$ is the PMF, $z_1$ and $z_2$ are points in aqueous solution on opposite sides of the membrane, $k_B$ is Boltzmann constant, and $T$ is the absolute temperature.

The distribution coefficient, log $D_{MW}$, was computed as:

$$logD_{\mathrm{MW}} = \log_{10}\left(\frac{P_0 10^{\mathrm{pH}-pK_a} + P_1}{10^{\mathrm{pH}-pK_a} + 1}\right) \quad (S2)$$

where $P_0$ is the partition coefficient of a neutral drug form, and $P_1$ is the partition coefficient of a charged (protonated) drug form, p$K_a$ is the acid dissociation constant for the drug (7.0 for dofetilide (40)). Using equation **S1**, we computed $K_0$ = 3.14±1.17, and $K_1$ = 0.48±0.35, and standard errors were estimated via propagation of uncertainties. The composite distribution coefficient was computed using equation **S2** to be log $D_{MW}$ = 0.32 ±0.13 at physiological pH = 7.2 used in this work and using literature p$K_a$ value of of 7.0 (40). Standard errors were estimated via propagation of uncertainties.

To compute the neutral drug translocation rate across membrane (7.96 ± 1.37 $ms^{-1}$), we used Kramer's transition rate approximation, as was done previously (41, 42) using Hummer's method (36) for computing diffusion coefficient profiles across the POPC membrane. The symmetrized diffusion coefficient profiles (main text **Figure 2D**) for charged and neutral dofetilide are shown from bulk aqueous solution ($z$=-40 Å) to membrane center ($z$=0 Å) and indicate a rapid 10-fold drop of diffusion coefficients for both charged and neutral and neutral dofetilide as the drug molecules start interacting with lipid membranes, similar to previous observations (43, 44). The local diffusion coefficient near the membrane center, used to estimate the crossing rate, was computed to be $\mathcal{D}_{\mathbf{M}}$ **=** 0.44 ± 0.15 x $10^{-6}$ $cm^2$ $s^{-1}$, and the curvatures of the binding well (0.083) and peak (-0.165), are estimated from second derivatives of second-order polynomial fits to the relevant portion of each respective free energy profile segment.

**hERG Open State Model Generation**

*Refinement of cryo-EM hERG structure:* The 3D coordinates of hERG (PDB: 5VA2) obtained via cryogenic electron microscopy (cryo-EM) were used as a template (45). This structure is mostly complete except for some extracellular loops; namely, residues 434-451 (between helices S1 and S2), 512-519 (between helices S3 and S4), and 578-582, 598-602 (in the pore loop region) are unresolved. The template was truncated beyond S668 on the S6 helix, eliminating intracellular PAS and C-terminal domains, leaving only the voltage sensing domain (VSD) and pore domain (PD) portions for our study (1,004 residues in total for the homotetramer). ROSETTA symmetry methods (46), and *de novo* loop modeling protocols (46-48) were used to generate the missing loop regions. In this process, 3- and 9-mer protein fragments from structures in the PDB were first obtained using the Robetta fragment server for the target hERG amino acid sequence (49). A conformational search using those structural fragments was performed according to cyclic coordinate descent (CCD) and kinematic loop closure (KLC) algorithms (50, 51). 20,000 initial poses were generated and the top 10% were filtered according to total energy score. This subset of poses were clustered (52) with a 2.0 Å root-mean-square deviation (RMSD) cutoff, and a low scoring structure (-984.884 Rosetta Energy Units (REU)) was selected from the most frequently sampled ensemble of models to undergo side chain relaxation (48, 53), in which protein backbone atoms are fixed while side chains are repacked using rotamer libraries. In this phase, simulated annealing followed by gradient minimization of the protein side chain torsional space was performed, and resultant poses were accepted or rejected based on the Metropolis criterion. 20,000 poses were again generated in this relaxation phase, the top 10% filtered and clustered, and from this subset a low scoring (-1961.173 REU or -1.95 REU/residue) structure was selected for use in the remainder of this study using Molecular Dynamics (MD) simulations. An initial structure of S641A hERG mutant for MD simulations was obtained using the wild-type channel model described above via side chain alteration of Ser 641 residues of four protein chains using UCSF Chimera.

**hERG Model Equilibration and Stability**

Our refined hERG model, generated with ROSETTA, was embedded in a POPC lipid bilayer and solvated with a 150mM aqueous KCl solution using CHARMM36 all-atom force fields and TIP3P water model using CHARMM-GUI web toolkit (23). Assembled systems consisted of ~128,000 atoms and were simulated with NAMD 2.12 (34) at a constant pressure of 1 atm and at a physiological temperature of 310 K. The systems were equilibrated for 90 ns using staged, extended equilibration methodology outlined in **Table S4** that we found to be imperative for maintaining protein stability. After 90 ns initial equilibration, which is a typical simulation time necessary to test membrane protein model structural stability and relieve steric clashes based on our previous ion channel simulations (54), the system was further simulated for 1 μs using Anton 2 software version 1.27.0 (i.e. achieving ~1.1 μs total simulation time). The channel pore remained well hydrated and open during this simulation time period, with the pore domain RMSD drifting up to ~3 Å and the SF RMSD remaining mostly within ~1 Å as can be seen in **Figure S5**. The HOLE program (55) was used to compute pore radius profiles.

**Potassium Ion Permeation Simulations for hERG and $K_V$1.2/2.1 chimera.**

*Voltage application protocol:* After 90 ns equilibration MD simulation described above, the equilibrated hERG model in the hydrated POPC bilayer was subsequently used for validating potassium ion permeation by applying transmembrane voltage during multi-microsecond simulations on Anton 2. To do so, a uniform electric field was applied in *z* direction, which can be computed as:

$$E_z = \frac{V}{L_z \cdot 43.5} \qquad \text{(S3)}$$

Where $V$ is the voltage in mV, $E_z$ is the $z$ component of the electric field vector in kcal/(mol·Å·e), and $L_z$ is the length of the unit cell in $z$ direction in Å. A factor of 43.5 was used to convert from mV/Å to kcal/(mol·Å·e). To prevent changes in applied voltage due to fluctuations in $L_z$, the system was simulated in the *NVT* ensemble. A voltage of +750 millivolts (mV) was selected as it was demonstrated to evoke fast ion permeation in models of $K_V$1.2/2.1 paddle chimera in a previous study (15). hERG simulations under applied +750 mV voltage were run for 5 μs with multiple $K^+$ conduction evens detected. Similar simulations were run for S641A hERG mutant, where no conduction events were observed during the ~2 μs simulation. Lipid membrane electroporation was not observed during these simulations. A smaller applied voltage of +500 mV resulted in only a single conduction event during our multi-microsecond simulations (data not shown) .

*$K_V$1.2/2.1 chimera ion conduction simulations:* The $K_V$1.2/2.1 chimera model was similarly evaluated for $K^+$ conduction through application of +750mV voltage in *NVT* Anton 2 simulation using the same voltage application protocol as defined above. We selected the $K_V$1.2/2.1 paddle chimera channel (Protein Data Bank ID: 2R9R; (56)) as our initial structure, as it has been successfully used in assessing $K^+$ ion conduction previously (15). The β subunit and tetramerization domains were then removed, as in that study. This truncated structure, henceforth referred to as the "$K_V$1.2/2.1 chimera" model, was then used to build an initial system with the CHARMM- GUI Membrane builder web toolkit (57). This system was then equilibrated for 50 ns in the *NPT* ensemble using NAMD and the same parameters and conditions as specified above in the general MD Simulation setup. In total, the system contained ~127,000 atoms, including 72 potassium ions, 64 chloride ions for charge equalization, and a lipid bilayer of 281 POPC lipids. The system was subsequently run in the *NVT* ensemble on Anton 2 for 5 μs. Voltage was applied using the protocol described above, but with the different electric field strength $E_z$ scaled to account for a new unit cell height $L_z$ and ensure the same transmembrane voltage of +750 mV.

*Counting permeating $K^+$ ions during channel conduction simulations:* Conduction for both the hERG and Kv1.2/2.1 Chimera models was evaluated using the same protocol, wherein ion *z*-positions with respect to channel selectivity filter (SF) backbone center of mass (COM) were recorded and flagged for their presence within the ion channel pore, and filtered for ion z-coordinate positions for plotting. In this protocol, the whole system was re-centered with respect to the COM of the channel. Then, for every ion and frame of the simulation, the *xy* component of the position vector of a particular ion was recorded as a radial distance from the *z* axis. These radii along with ion *z* positions were then used to determine individual ion localization with respect to the channel pore, per ion and per frame: when this *xy* component was less than 10 Å, a distance inclusive to the entirety of the channel's conductive pathway, and the $z$-coordinate of the ion resided within $z$ bounds encompassing the entire channel, then the particular z-coordinate position was flagged as residing in the pore. The bounds used were as follows: hERG: -35≤ $z$ ≤ 30 Å; for $K_V$1.2/2.1 Chimera: -16 ≤ $z$ ≤ 20 Å. Only flagged z-positions were plotted in $K^+$ conduction time series, and permeation events were counted from the resultant plots.

*Pore hydration measurements*: To measure pore hydration, a VMD script was used to record the quantity of water molecules occupying the pore cavity of a given ion channel. The pore cavity was defined as the region of the ion channel from the bottom of the S6 helix to the base of the selectivity filter, incorporating the "cavity" ("S5") ion position formed by the hydroxyls of Ser626 of hERG or Thr370 for $K_V$1.2/2.1 Chimera. Water molecules were counted if they were within a cylindrical selection defined to include the whole pore cavity. This cylinder has a radius of 10 Å, a base parallel to the membrane, and a height equal to the distance between the selectivity filter and the bottom of the S6 helix. Prior to measuring pore hydration, the entire system was re-centered about the center of mass of the selectivity filter.

*Selectivity filter residue $\varphi$ and $\psi$ measurements:* Protein backbone torsional angles ($\varphi$ and $\psi$) were measured for the selectivity filter residues SVGFG (hERG) or TVGYG ($K_V$1.2/2.1 Chimera) using VMD's Timeline plugin.

**hERG Drug "Flooding" Simulations**

Multi-microsecond unbiased drug "flooding" MD simulations were used to study dofetilide binding pathways to hERG. They were initiated from building using CHARMM-GUI and equilibrating using NAMD channel/membrane systems solvated in 0.15 M KCl aqueous solutions containing 0.025 M (i.e., 25 mM) of each drug. Initially, drug molecules were randomly placed around the protein with the CHARMM script excluding lipid membrane and the channel itself and also ensuring optimal distances between adjacent molecules. Then, the hydrated lipid membrane was added using CHARMM-GUI, and the entire system was equilibrated for 90 ns with NAMD in the *NPT* ensemble at 310 K and 1 atm pressure using the same protocol as for drug-free hERG simulations described above (and highlighted in **Table S4**). The system sizes and compositions are also similar to ones used in our drug-free hERG simulations discussed above. The equilibrated systems were further simulated on Anton 2 in the *NPT* ensemble for 2.5 μs for both DOFC and DOFN containing systems. No external voltage or restraints were applied in those simulations. These simulations are similar to ones used in our previous study (37), where we determined drug binding to voltage-gated $Na^+$ channel $Na_VAb$ through both a traditional aqueous pore binding pathway, and through lipid-facing channel openings, or fenestrations. Here, we also aimed to explore both aqueous and lipid-mediated pathways for dofetilide binding to hERG.

**Umbrella sampling MD simulations to compute hERG-Drug Binding Energetics and Kinetics.**

*Setup for US MD simulations using steered MD:* In order to ensure sufficiently randomized starting orientations of dofetilide with which to seed each US window, we performed a set of steered molecular dynamics simulations, for both charged and neutral dofetilide and the open hERG channel. Using CHARMM-GUI, CHARMM36 all-atom force fields and TIP3P water model, each hERG model was embedded in a POPC lipid bilayer and solvated with a 150 mM aqueous KCl solution. Assembled systems consisted of ~132k atoms and were simulated with NAMD 2.12 in the *NPT* ensemble at a constant pressure of 1 atm and at a physiological temperature of 310 K. These simulations began with an initial equilibration run in which the drug was harmonically restrained at $z$=-50 Å (with respect to the SF COM), i.e. at the intracellular side of the channel, below the pore (see e.g. main text **Figure 6A**), but allowed to rotate freely. After equilibrating for ~50ns, 5 different initial configurations were chosen to begin steered molecular dynamics runs, in which the drug was pulled, in 0.5 Å increments for each 1 ns of the simulations, at constant applied force, from bulk water ($z$=-50 Å) into the pore, $z$=-5 Å, i.e. to a point 5 Å below the COM of selectivity filter. Total simulation time for each pulling simulations was 91 ns. From these trajectories, initial umbrella sampling windows were seeded with each window randomly chosen for a respective drug $z$ position from the 5 pulling simulations and shuffled such that no two neighboring US windows were chosen from the same simulation. In this way, every window in the umbrella sampling simulations would begin with a sufficiently randomized drug orientation, with neighboring windows dissimilar from each other. This way, for hERG - drug US MD simulations 91 independent US windows were seeded, spaced in 0.5 Å intervals from −50 Å to -5 Å with respect to COM of hERG selectivity filter $C_\alpha$ atoms hERG SF. The pore $C_\alpha$ atoms and all SF backbone non-H atoms were subject to 1.0 kcal/mol/Å$^2$ harmonic restraints during initial equilibration and pulling runs. They were gradually decreased to 0.2 kcal/mol/Å$^2$ during the first 8 ns of each US MD run, as described in more detail below.

*US simulations for dofetilide translocation across the hERG pore:* These US simulations were performed in roughly the same way as for drug-membrane partitioning simulations described above, but with drug center of mass held with respect to selectivity filter $C_\alpha$ atoms, instead of membrane COM as described above. Such an approach was used previously for a similar system (58). For these US simulations, dofetilide COM position was spaced at 0.5 Å intervals from $z$= -5 Å to $z$ = -50 Å with respect to COM of the selectivity filter $C_\alpha$ atoms, resulting in 91 US windows in total for each charged and neutral forms of the drug. The initial US MD simulation systems were prepared as described above. For each window, a force constant of 10.0 kcal/mol/Å$^2$ was applied to harmonically restrain the drug $z$ position within each window, and an additional 5 kcal/mol/Å$^2$ cylindrical restraint was applied in order to prevent drift of the drug in the $xy$ plane using the collective variables (COLVAR) functionality of NAMD. The first set of the US simulations were run without restraints on the hERG open model, but in this case for some US windows the pore was found to be closing and/or SF distorted. Since there were substantial instabilities in unrestrained runs of hERG in the open state, we additionally ran the same set of simulations, but with moderate restraints on the hERG pore $C_\alpha$ atoms, staged down from 1kcal/mol/Å$^2$ to 0.2kcal/mol/Å$^2$ over the course of the initial 8 ns of each US window, which ran for 40 ns total. The PMF profiles were computed using WHAM, with the initial 10ns of each window discarded to account for equilibration. Error bars were computed from standard error of the mean of 5 ns blocks averaging PMF data from simulations times from 10 to 40ns. The effective equilibrium dissociation constant, $K_{D}$, was computed from the PMF profiles, as done previously (58).

To compute the drug-channel association or ingress "on" rate constant for dofetilide, we assumed that the binding process is a purely diffusion-limited reaction, and that the reaction (drug binding) is fast once the molecule enters the reactive region. Diffusion coefficient, $\mathcal{D}(z)$, profiles for charged and neutral dofetilide, going from the intracellular bulk into the hERG pore were obtained using Laplace transform of drug position autocorrelation function (36) as was done in our recent studies on drug - membrane partitioning (37, 38). Having both the diffusion coefficient profile, and PMF along the reaction coordinate, we computed the ingress rate constant using a formulation of the Debye-Smoluchowski equation (59, 60), previously implemented in an ion permeability study (61),

$$k_{on}{}^{-1} = \frac{1}{\pi R^2} \int_{z_{out}}^{z_{bind}} e^{\frac{W(z)}{k_B T}} \mathcal{D}(z)^{-1} dz \quad \text{(S4)}$$

where $R$ designates the radius of a cylinder (10 Å) that encompasses the hERG channel pore, and $\mathcal{D}(z)$ is the local diffusion coefficient along the reaction coordinate. Having already computed $K_{\mathrm{D}}$ and $k_{\mathrm{on}}$, we can trivially compute drug-channel dissociation or "off" rate constant $k_{\mathrm{off}}$ since $K_{\mathrm{D}} = k_{\mathrm{off}} / k_{\mathrm{on}}$ (62), so

$$k_{\mathrm{off}} = k_{\mathrm{on}} \cdot K_{\mathrm{D}} \quad \text{(S5)}.$$

#### SI Results and Discussion

##### Details of hERG Model K+ Conduction under the Applied Voltage

Ten $K^+$ conduction events were observed in the wild-type hERG simulations, as outlined in the **Figure S6.** Seven of these events occur between $t$ = 300 ns to $t$ = 700 ns and are shown in more detail in the main text **Figure 3**. A $K^+$ ion that resides in the pore cavity has a $z$ coordinate (with respect to the SF backbone center of mass) $-25 \leq z \leq 4$ Å. $K^+$ ions within the SF predominately occupied the [S1 S3] state, corresponding to $z$ = -4 Å and $z$ = +4 Å coordinates respectively. The [S2 S4] state occurs transiently and only coincidentally with the escape of an S1 ion. Both direct ("hard") and water-mediated ("soft") knock-on conduction events were observed. $K^+$ conduction ceased after 1.75 μs, coinciding with a marked reduction in the channel pore diameter and consequently pore dehydration (**SI Figure S7C).** Conduction events are preceded and accompanied by marked changes in the SF conformation as explored in **Figure S7A and S7B**. In particular, during the 300 ns conduction interval, the G628 and F627 residues of the selectivity filter adopt a flipped confirmation in which their carbonyl oxygens coordinate with water molecules located behind the selectivity filter, as opposed to those in the S1 and/or S0 sites. This consequently widens the top of selectivity filter, preventing occupancy of the S0 site and potentially facilitating escape of $K^+$ from S1. As a result, ion permeation transitions predominantly from the "cavity, S3, and S1 state" to "cavity and S3 state" throughout the simulation.

Conduction temporarily ceases after $t$ = 700 ns, coinciding with the elimination of the S1 occupancy site and pinching at the S2 site in the selectivity filter, which is measured as the respective increase in opposing chain $C_\alpha$ distances for Gly628 and decrease in $C_\alpha$ distances for Gly626 (see **Figure S7B,** green and blue traces). The three remaining conduction events occur from $t$ = 1400 ns to $t$ = 1700 ns and happen during closure of the pore at the intracellular gate, as indicated by pore dehydrating at $t$ = 1400 ns (See **Figure S7C**). For the two periods of conduction, the $\varphi$ and $\psi$ backbone dihedral angles of Gly628, the top-most selectivity filter residue, exhibit 60 to 120-degree fluctuations and adopt asymmetrical conformations (see **Figure S7A**). In contrast, the non-conducting portion is intimated by a 120-degree shift in Gly628 $\varphi$ and $\psi$ angles from both the preceding and anteceding conduction segments and reflects the elimination of the S1 site.

The manner in which conduction ceases and resumes prior to pore collapse is of note. Restoration of the conductive selectivity filter state at $t$=1400 ns is mediated by subsequent knock-on events, the first of which results in an [S3 S4] ion arrangement **(Figure S6A).** This arrangement widens the S2 site to make it accessible to water. Once S2 is solvated from the extracellular side, a subsequent knock-on attempt induces the SF blocking $K^+$ ion at S3 site to transition to S1, but not before partially exiting the SF to exchange its S2 position with an extracellular water. This reshuffling of the SF blocking ion across collapsed region of the SF and the reintroduction of water to S2 induces rotations in Gly626 and Phe627, resulting in a state that resembles the original conducting orientation, and conduction resumes briefly. However, the pore cavity narrows substantially by 1.4 μs, as measured by pore solvation and is no longer readily accessible to potassium ions, and further conduction ceases (**Figure**

**S7C**). The pore radius at the intracellular gate is substantially reduced beyond this point ( **Figure S6A**) indicating channel pore hydrophobic collapse.

**Details of $K_V$1.2/2.1 Chimera Model $K^+$ Conduction under the Applied Voltage**

The $K_V$1.2/2.1 chimera model was similarly induced to conduct by applying +750mV. **Figure S6B** presents the entire 5.05 μs ion conduction trajectory in the $K_V$1.2/2.1 chimera model. The model conducts early into simulation: from 50-350 ns, 9 $K^+$ ions permeate, adopting either the [S1 S3] or [S2 S3] states.

The pore of the $K_V$1.2/2.1model remains open for a longer duration than the hERG model pore and avoids hydrophobic collapse, as indicated by their respective pore radii profiles produced by HOLE program (**Figure S6B**). However, the putative $K^+$ ion position S0-S4 SF sites undergo substantial disordering by the end of the simulation. This is reflected in the SF collapse, which occurs by 4.3 μs, resulting in a ~0.8 μs of ultra-rapid uncontrolled conduction. This period is visually apparent in the backbone torsional angles $\varphi$ and $\psi$ plots of the selectivity filter residues in **SI Figure S8A**. This collapsed state was preceded by an asymmetrical increase in Thr370 $C_\alpha$-$C_\alpha$ by 1 to 3 angstroms, and aggressive pinching in Gly372, the center of the selectivity filter. These factors define the locked state in which ions are located solely at S4 site from 3.2 – 4.3 μs. One factor to consider is what role the rapid de-hydration of the pore cavity at *t*=2.9 μs has in establishing the conditions that facilitated pore collapse during the last portion of the simulation.

Prior to this collapse, there is a longer period of non-conductivity. This period begins with selectivity filter de-hydration and pinching at Gly374 starting at about 400 ns (**Figure S7B**, green traces). However, the ions are locked in the [S2 S3] state, in which conduction predominantly occurred earlier in the simulation (**Figure S6B**). Examining the intracellular gate portion of the model in **Figure S6B** indicates occasional ion visitations that should present opportunities for knock-on events. However, there is also a relative reduction in pore hydration **(Figure S8C**) in comparison to the initial conducting phase not reflected in the pore radii measurements that may contribute to non-conduction. The $K_V$1.2/2.1 model has fewer fluctuations in SF $\varphi$ and $\psi$ angles and $C_\alpha$-$C_\alpha$ distances than in hERG during their respective periods of conduction, suggesting that the pore is simply temporarily and partially closing for $K_V1.2/2.1$ model. However, whether the gross structural features of the states of pore partial dehydration and changes in a SF geometry represent a particular ion channel state remain unclear and would require a separate study beyond the scope of this work.

**Molecular simulation model limitations**

In this study we used a prototypical single component POPC lipid membrane, which has been used in numerous previous MD simulation studies of hERG and other ion channels. Lipid membrane of cardiomyocytes has a much more complex composition and in addition to phosphatidylcholine includes phosphatidylethanolamine, anionic phospholipids, sphingolipids and cholesterol to name a few (63, 64). hERG activity is also known to be modulated by these and other membrane components such as PIP2 (65), polyunsaturated fatty acids (66), ceramides (67) and others, as was recently reviewed in (68). However, using a single-component lipid membrane in our simulations is warranted by absence of lipophilic pathway for dofetilide hERG binding, which might have been influenced by lipid membrane composition. Moreover, usage of multi-component lipid membranes will necessitate the use of much longer simulations to sample lipid diffusion in the vicinity of the protein and achieve converged results. Thus, the usage of one-component lipid membrane should suffice for the purpose of the work described here, the development of a robust proof-of-principle platform for multi-scale testing of drug cardiac pharmacology.

It should be noted, however, that some quantities like drug water-membrane partitioning coefficients can be significantly influenced by lipid membrane composition. This can possibly explain why our computed $\log D_{\mathrm{MW}}$ value of 0.32 ± 0.13 is quite different from that of 2.08, derived from a high performance liquid chromatography (HPLC) experiment of dofetilide partitioning from water into an Immobilized Artificial Membrane (IAM) (69). Besides different membrane structure and chemical composition, it is also not clear if we can make a direct quantitative comparison between drug partitioning results of a small lipid bilayer patch in MD simulations (70) and those macroscopic experimental data, which also employ empirical relations to estimate $\log D$ in IAM experiments (71).

For dofetilide – hERG simulation system, we considered drug binding only to an open conducting channel state, for which we have structural data. However, experimental data suggest that dofetilide binds considerably stronger to inactivated hERG state, with $IC_{50}$ in the low nanomolar range (72-76). To include dofetilide binding to the inactivated state into our functional model for this prototype study, we used the experimental ratio of binding affinities to these two states (see the companion paper, part 2). However, future model development will necessitate a structural model of the channel inactivated state. We did attempt to build an inactivated state hERG model using previously suggested conformational restraints: N629···S620 intra-subunit hydrogen bonds (similar to E71···D80 interactions in KcsA) (77). This resulted in a SF constriction and widening of putative drug binding pockets under the SF, all indicative of a possible inactivated state. However, those hydrogen bonding interactions were found to be broken in μs-long unbiased MD simulations (data not shown), restoring typical hERG behavior observed in simulations presented in this work. However, we also found that ~2 μs-long simulations of a S641A hERG mutant (based on the WT structure) resulted in a non-conductive channel PD structure, with the SF pinched at the top (at the level of G628 residues). We saw similar, but more transient SF pinching events in our $K_V$1.2/2.1 and WT hERG conduction runs, which resulted in prolonged loss of conduction. The S641A mutation is known to facilitate hERG inactivation, but it also practically eliminates channel block by E4031, a close analog of dofetilide (78). Thus, we could not use this mutant channel structural model to probe dofetilide binding to the inactivated state, and we are still in search of inactivated hERG structural models with high affinity for dofetilide and/or other hERG blockers.

#### SI Tables

**Table S1. Partial atomic charges for charged (DOFC) and neutral (DOFN) dofetilide models.**

(Optimized charge values are shown by asterisk)

| Drug | Total Charge | Drug | Total Charge |
|---|---|---|---|
| **DOFC** | 1.000 | **DOFN** | 0.000 |
| C1 | 0.149 | C1* | -0.392 |
| N1* | -0.006 | N1* | -0.401 |
| C2* | -0.495 | C2* | 0.240 |
| C3* | -0.250 | C3* | -0.559 |
| C4* | 0.310 | C4* | 0.296 |
| C5 | -0.115 | C5 | -0.115 |
| C6 | -0.111 | C6 | -0.111 |
| C7 | 0.219 | C7 | 0.219 |
| C8 | -0.111 | C8 | -0.111 |
| C9 | -0.115 | C9 | -0.115 |
| N2 | -0.466 | N2 | -0.466 |
| S1 | 0.439 | S1 | 0.439 |
| O1 | -0.384 | O1 | -0.384 |
| O2 | -0.384 | O2 | -0.384 |
| C10 | -0.019 | C10 | -0.019 |
| C11 | 0.162 | C11* | 0.010 |
| C12 | -0.014 | C12 | -0.021 |
| O3 | -0.391 | O3 | -0.39 |
| C13 | 0.219 | C13 | 0.219 |
| C14 | -0.113 | C14 | -0.113 |
| C15 | -0.115 | C15 | -0.115 |
| C16 | 0.219 | C16 | 0.219 |
| C17 | -0.115 | C17 | -0.115 |
| C18 | -0.113 | C18 | -0.113 |
| N3 | -0.466 | N3 | -0.466 |
| S2 | 0.439 | S2 | 0.439 |
| O4 | -0.384 | O4 | -0.384 |
| O5 | -0.384 | O5 | -0.384 |
| C19 | -0.019 | C19 | -0.019 |
| H1 | 0.090 | H1 | 0.090 |
| H2 | 0.090 | H2 | 0.090 |
| H3 | 0.090 | H3 | 0.090 |
| H4 | 0.090 | H4 | 0.090 |
| H5 | 0.090 | H5 | 0.090 |
| H6 | 0.090 | H6 | 0.090 |
| H7 | 0.090 | H7 | 0.090 |
| H8 | 0.115 | H8 | 0.115 |
| H9 | 0.115 | H9 | 0.115 |
| H10 | 0.115 | H10 | 0.115 |
| H11 | 0.115 | H11 | 0.115 |
| H12 | 0.323 | H12 | 0.323 |
| H13 | 0.090 | H13 | 0.090 |
| H14 | 0.090 | H14 | 0.090 |
| H15 | 0.090 | H15 | 0.090 |
| H16 | 0.090 | H16 | 0.090 |
| H17 | 0.090 | H17 | 0.090 |
| H18 | 0.090 | H18 | 0.090 |
| H19 | 0.090 | H19 | 0.090 |
| H20 | 0.115 | H20 | 0.115 |
| H21 | 0.115 | H21 | 0.115 |
| H22 | 0.115 | H22 | 0.115 |
| H23 | 0.115 | H23 | 0.115 |
| H24 | 0.323 | H24 | 0.323 |
| H25 | 0.090 | H25 | 0.090 |
| H26 | 0.090 | H26 | 0.090 |
| H27 | 0.090 | H27 | 0.090 |
| HN | 0.318 | | |

#### Table S2. Gas-phase neutral dofetilide (DOFN) – water interactions.

| | Water interaction distance (A) | | | Water interaction energy (kcal/mol) | | |
|---|---|---|---|---|---|---|
| | QM | MM | *Difference* | QM | MM | *Difference* |
| N1 | 3.077 | 3.13 | *0.05* | -5.839 | -6.017 | *-0.178* |
| N2 | 3.205 | 3.31 | *0.10* | -3.572 | -2.582 | *0.990* |
| N3 | 3.26 | 3.21 | *-0.05* | -4.137 | -5.059 | *-0.922* |
| O1 | 3.003 | 2.95 | *-0.05* | -6.363 | -6.377 | *-0.014* |
| O1 | 2.986 | 2.94 | *-0.05* | -6.34 | -6.396 | *-0.056* |
| O1 | 3.049 | 2.95 | *-0.10* | -5.384 | -5.409 | *-0.025* |
| O2 | 3.053 | 2.95 | *-0.10* | -6.019 | -6.677 | *-0.658* |
| O2 | 2.977 | 2.93 | *-0.05* | -6.822 | -7.164 | *-0.342* |
| O2 | 3.065 | 2.97 | *-0.10* | -5.515 | -5.753 | *-0.238* |
| O3 | 3.01 | 2.91 | *-0.10* | -4.847 | -5.452 | *-0.605* |
| O4 | 3.039 | 2.99 | *-0.05* | -5.764 | -6.268 | *-0.504* |
| O4 | 2.987 | 2.94 | *-0.05* | -6.254 | -6.867 | *-0.613* |
| O4 | 3.018 | 2.97 | *-0.05* | -5.721 | -6.023 | *-0.302* |
| O5 | 3.045 | 3.00 | *-0.05* | -5.844 | -6.039 | *-0.195* |
| O5 | 2.974 | 2.92 | *-0.05* | -6.761 | -6.833 | *-0.072* |
| O5 | 3.044 | 2.94 | *-0.10* | -5.713 | -5.814 | *-0.101* |
| H8 | 2.515 | 2.82 | *0.30* | -2.324 | -2.02 | *0.304* |
| H9 | 3.28 | 3.68 | *0.40* | -3.321 | -2.796 | *0.525* |
| H10 | 2.375 | 2.78 | *0.40* | -3.877 | -3.12 | *0.757* |
| H11 | 2.504 | 2.80 | *0.30* | -3.24 | -2.828 | *0.412* |
| H12 | 1.989 | 2.04 | *0.05* | -7.541 | -6.935 | *0.606* |
| H20 | 2.512 | 2.86 | *0.35* | -1.634 | -0.797 | *0.837* |
| H21 | 2.538 | 2.94 | *0.40* | -0.309 | 0.884 | *1.193* |
| H22 | 2.43 | 2.78 | *0.35* | -4.092 | -3.323 | *0.769* |
| H24 | 2.012 | 2.01 | *0.00* | -7.342 | -6.946 | *0.396* |
| ***RMSE*** | | | ***0.211*** | | | ***0.527*** |

#### Table S3. Gas-phase charged dofetilide (DOFC) – water interactions.

| | Water interaction distance (A) | | | Water interaction energy (kcal/mol) | | |
|---|---|---|---|---|---|---|
| | QM | MM | *Difference* | QM | MM | *Difference* |
| N2 | 3.237 | 3.29 | *0.05* | -2.26 | -1.975 | *0.285* |
| N3 | 3.148 | 3.20 | *0.05* | -3.155 | -2.233 | *0.922* |
| O1 | 3.086 | 2.99 | *-0.10* | -8.934 | -9.013 | *-0.079* |
| O1 | 2.973 | 2.92 | *-0.05* | -6.092 | -6.75 | *-0.658* |
| O1 | 3.067 | 2.97 | *-0.10* | -7.698 | -7.875 | *-0.177* |
| O2 | 3.037 | 2.99 | *-0.05* | -6.907 | -7.176 | *-0.269* |
| O2 | 3.006 | 2.96 | *-0.05* | -5.22 | -5.922 | *-0.702* |
| O2 | 3.081 | 2.98 | *-0.10* | -4.433 | -5.068 | *-0.635* |
| O3 | 3.114 | 2.91 | *-0.20* | -1.865 | -3.155 | *-1.290* |
| O4 | 7.296 | 7.30 | *0.00* | -0.851 | -0.618 | *0.233* |
| O5 | 4.481 | 4.88 | *0.40* | -3.364 | -1.704 | *1.660* |
| O5 | 3.04 | 2.94 | *-0.10* | -3.51 | -4.124 | *-0.614* |
| O5 | 3.092 | 2.94 | *-0.15* | -3.383 | -4.898 | *-1.515* |
| H8 | 2.529 | 2.83 | *0.30* | -5.074 | -4.223 | *0.851* |
| H9 | 2.839 | 3.24 | *0.40* | -5.555 | -3.318 | *2.237* |
| H10 | 2.356 | 2.76 | *0.40* | -5.84 | -4.289 | *1.551* |
| H12 | 1.981 | 2.03 | *0.05* | -9.487 | -7.887 | *1.600* |
| H20 | 2.331 | 2.73 | *0.40* | -6.588 | -4.648 | *1.940* |
| H21 | 2.356 | 2.76 | *0.40* | -6.025 | -3.831 | *2.194* |
| H22 | 2.207 | 2.61 | *0.40* | -7.815 | -5.072 | *2.743* |
| H23 | 2.298 | 2.70 | *0.40* | -7.732 | -5.814 | *1.918* |
| H24 | 1.943 | 1.99 | *0.05* | -12.94 | -10.529 | *2.411* |
| ***RMSE*** | | | ***0.246*** | | | ***1.443*** |

#### Table S4. Restraint regime for hERG MD equilibration simulations

| **Time, ns** | **Restraint, Kcal/mol/Å$^2$** | **Protein Domain** |
|---|---|---|
| **1-5** | **1.0** | **Backbone** |
| **5-10** | **1.0** | **Pore Domain** |
| **10-15** | **0.5** | **Pore Domain** |
| **15-20** | **0.25** | **Pore Domain** |
| **20-30** | **0.1** | **Pore Domain** |
| **30-40** | **0.1** | **Selectivity Filter** |

**SI Figures**

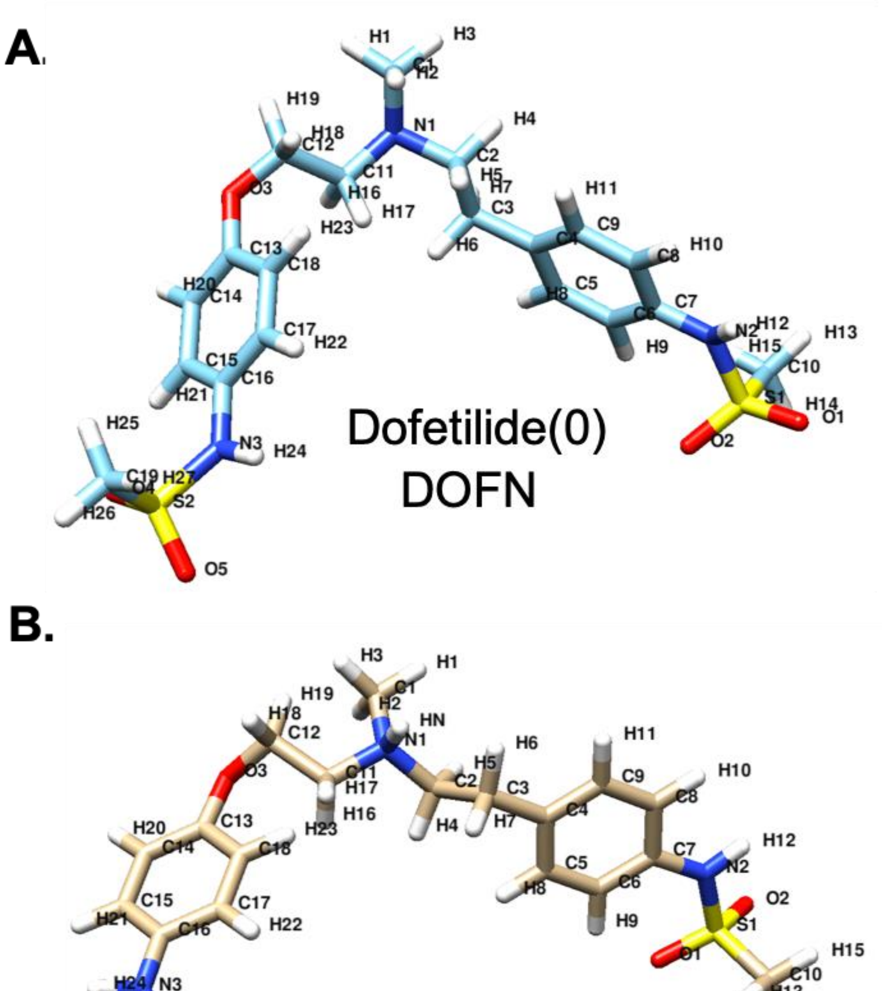

**Figure S1. Atom naming for charged and neutral dofetilide molecules. A.** DOFN or Dofetilide(0) –neutral dofetilide, **B.** DOFC or Dofetilide(+) – charged dofetilide

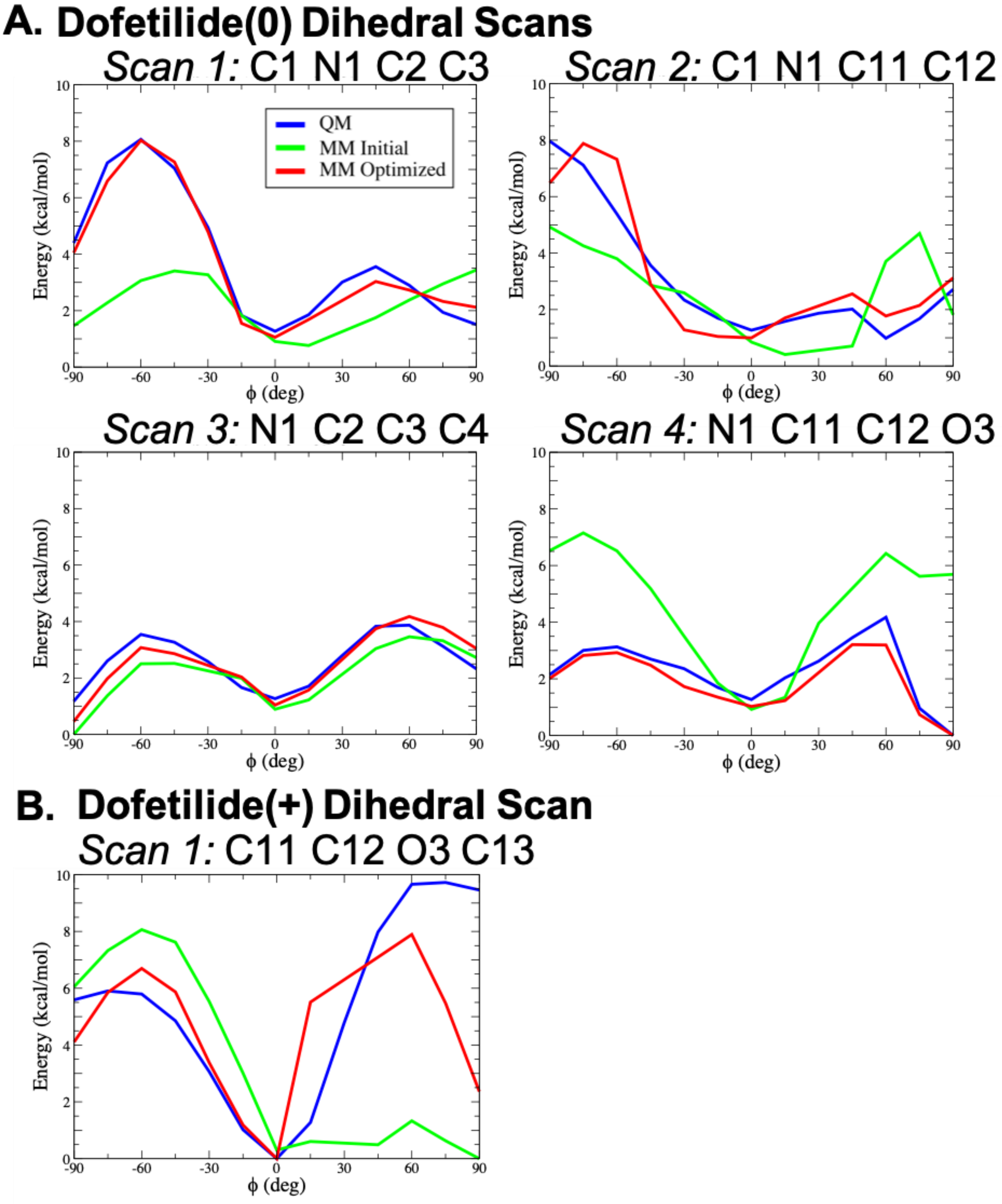

**Figure S2. Gas-phase torsional energy profiles for dofetilide models from quantum mechanical (QM), initial and optimized molecular mechanics (MM) calculations.** Atom names corresponding to the ones in Figure S1A and Figure S1B for (**A)** Dofetilide(0) or DOFN, and (**B)** Dofetilide(+) **or** DOFC, respectively, with corresponding topology and parameter files in the **Appendix SA1**.

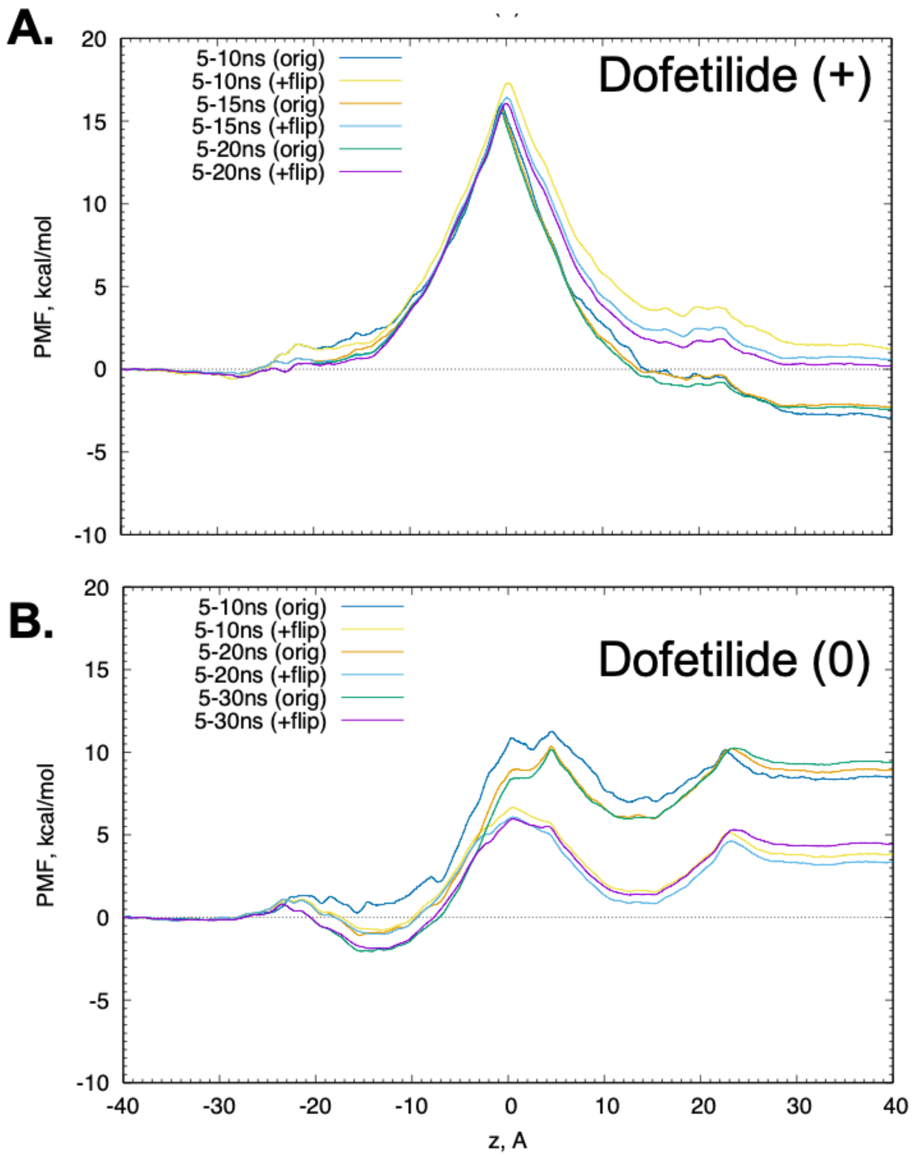

**Figure S3. Non-symmetrized free energy or potential of mean force (PMF) profiles for charged (A) and neutral (B) dofetilide crossing a POPC lipid bilayer computed from umbrella sampling MD simulations.** They were computed using weighted histogram analysis method (WHAM) discarding first 5 ns for each umbrella sampling window. In some cases, additional simulations with initial drug orientation rotated around the *z* axis (indicated as +flip) were performed to improve drug re-orientation sampling.

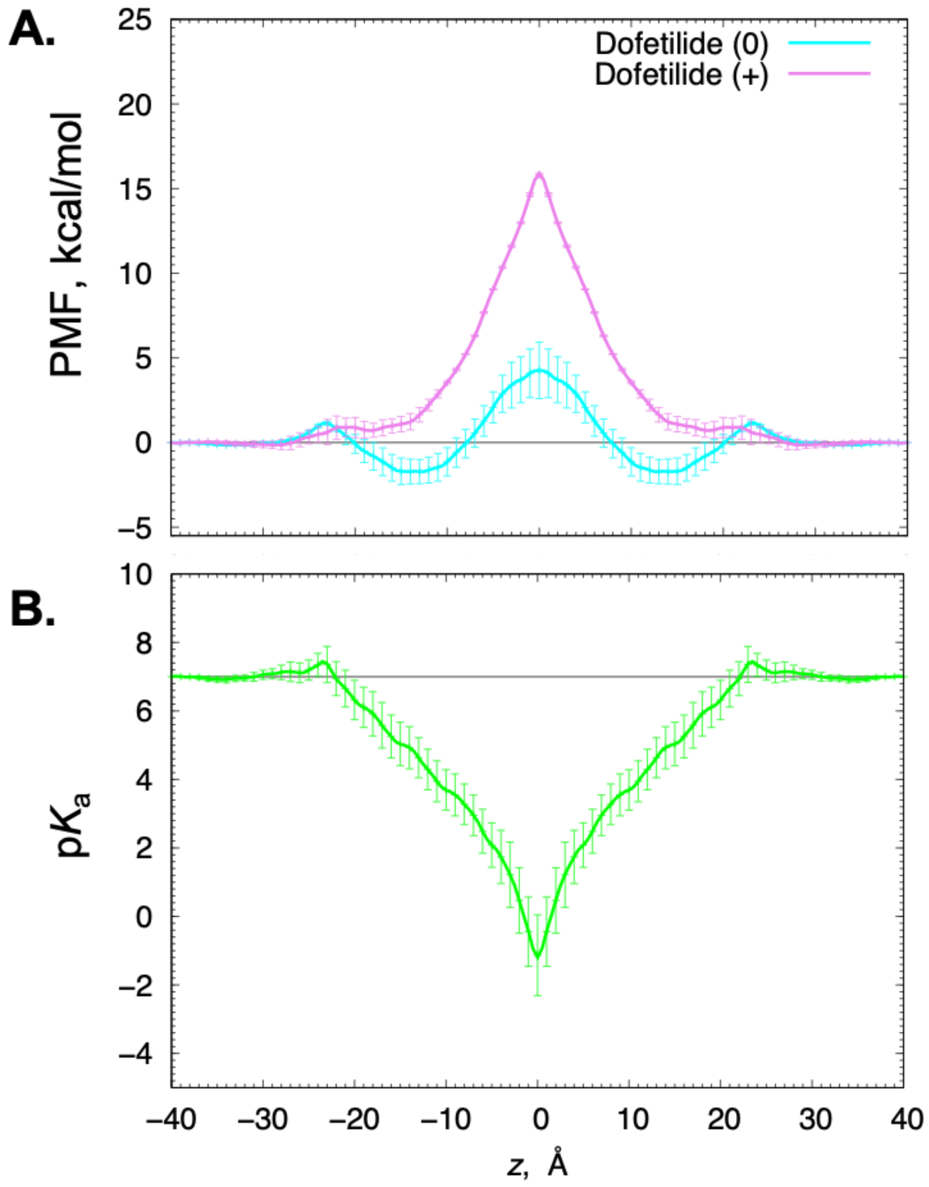

**Figure S4. Potential of mean force (PMF) and corresponding p$K_a$ profiles for dofetilide crossing a POPC lipid bilayer. (A)** Symmetrized PMF profiles of neutral (cyan) and charged (magenta) dofetilide. **(B)** Symmetrized p$K_a$ profile computed from PMFs in panel **A.** Aqueous p$K_a$ = 7.0 is shown as a horizontal black line. Error bars are computed as a measure of asymmetry.

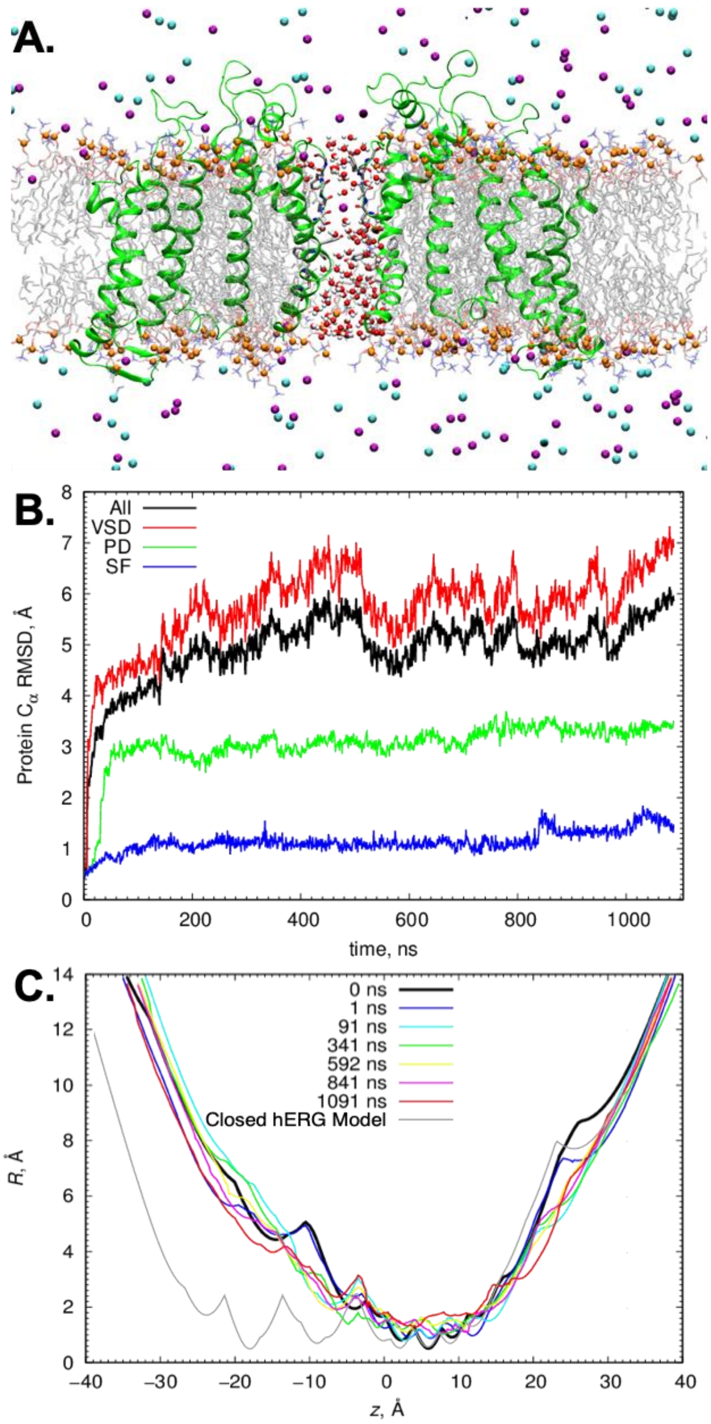

**Figure S5. Stability of open state hERG model. (A)** Molecular system snapshot at the end of 1 μs unbiased MD simulation. Two opposing hERG subunits are shown. Protein backbone is shown as green ribbons (with SF S624-G628, S6 helix Y652 and F656 residues shown as sticks), lipid headgroups as orange balls, lipid tails as grey think sticks, SF atoms shown as colored sticks, $K^+$ ions as purple and $Cl^-$ ions as cyan balls, pore waters as red/white spheres. Bulk water is not shown for clarity. (**B**) Protein $C_\alpha$ Root-mean-squared-deviation (RMSD) time series broken down by domain, and (**C**) pore radius profiles at selected time points**.**

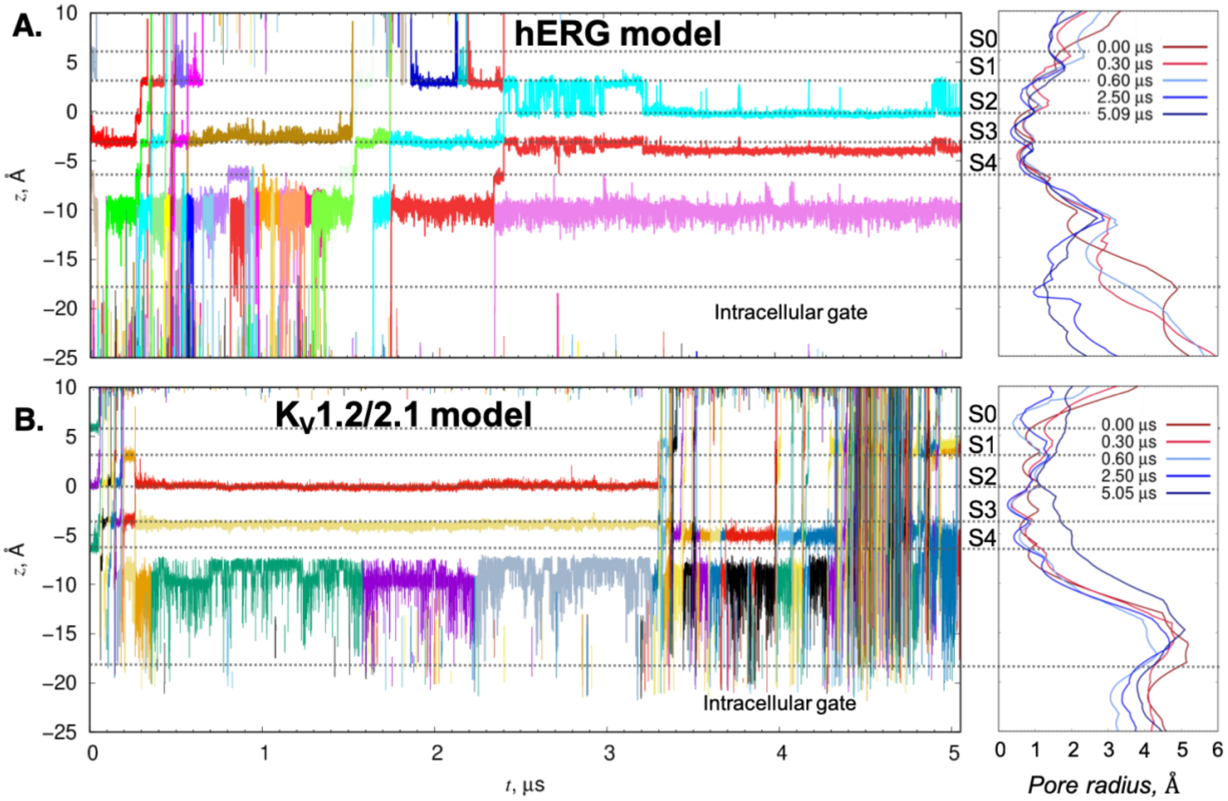

**Figure S6. Complete time series of K⁺ pore positions in hERG (A) and Kv1.2/2.1 chimera (B) channel models under an applied 750 mV voltage and corresponding pore radius profiles** *Left*: Individual $K^+$ *z*-coordinate positions (colored traces) for the full duration of the ~5 µs simulations. z-coordinate positions of SF $K^+$ ion occupancy sites are denoted with canonical S0-S4 notation, as is the location of the intracellular gate. *Right:* pore radii of the channel models measured at 5 timepoints to reflect pore narrowing and/or elimination of SF ion occupancy sites.

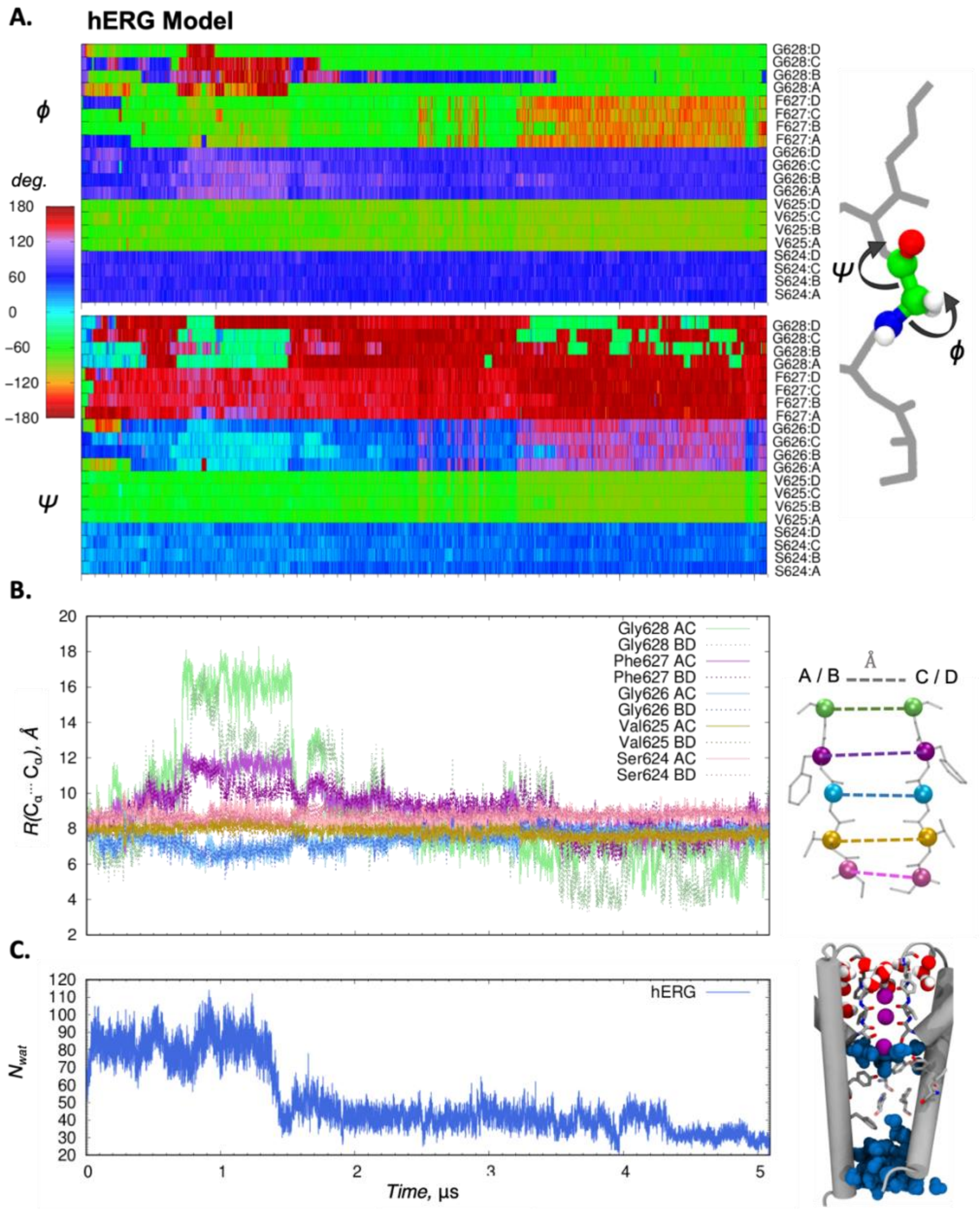

**Figure S7. Structural analysis of hERG from MD simulations under applied 750 mV voltage. (A)** *Left:* Time series of $\varphi$ and $\psi$ protein backbone dihedral angles of SF residues SVGFG. *Right:* Graphical representation of $\varphi$ and $\psi$ dihedral angles in the SF. **(B)** *Left*: Time series of distances between protein backbone $C_\alpha$ atoms between SF residues in the opposing chains. *Right:* Depiction of the distances being measured, with protein chains “A” and “B” opposite to “C” and “D”, respectively. **(C)** *Left:* Time series of pore hydration as measured by number of water molecules within the pore. *Right:* Snapshot of pore hydration at the final simulation frame ($t$ = 5.090 μs) . Relevant water molecules occupying pore are colored blue and correspond to the plotted quantity on the left. Water molecules behind SF are red/white.

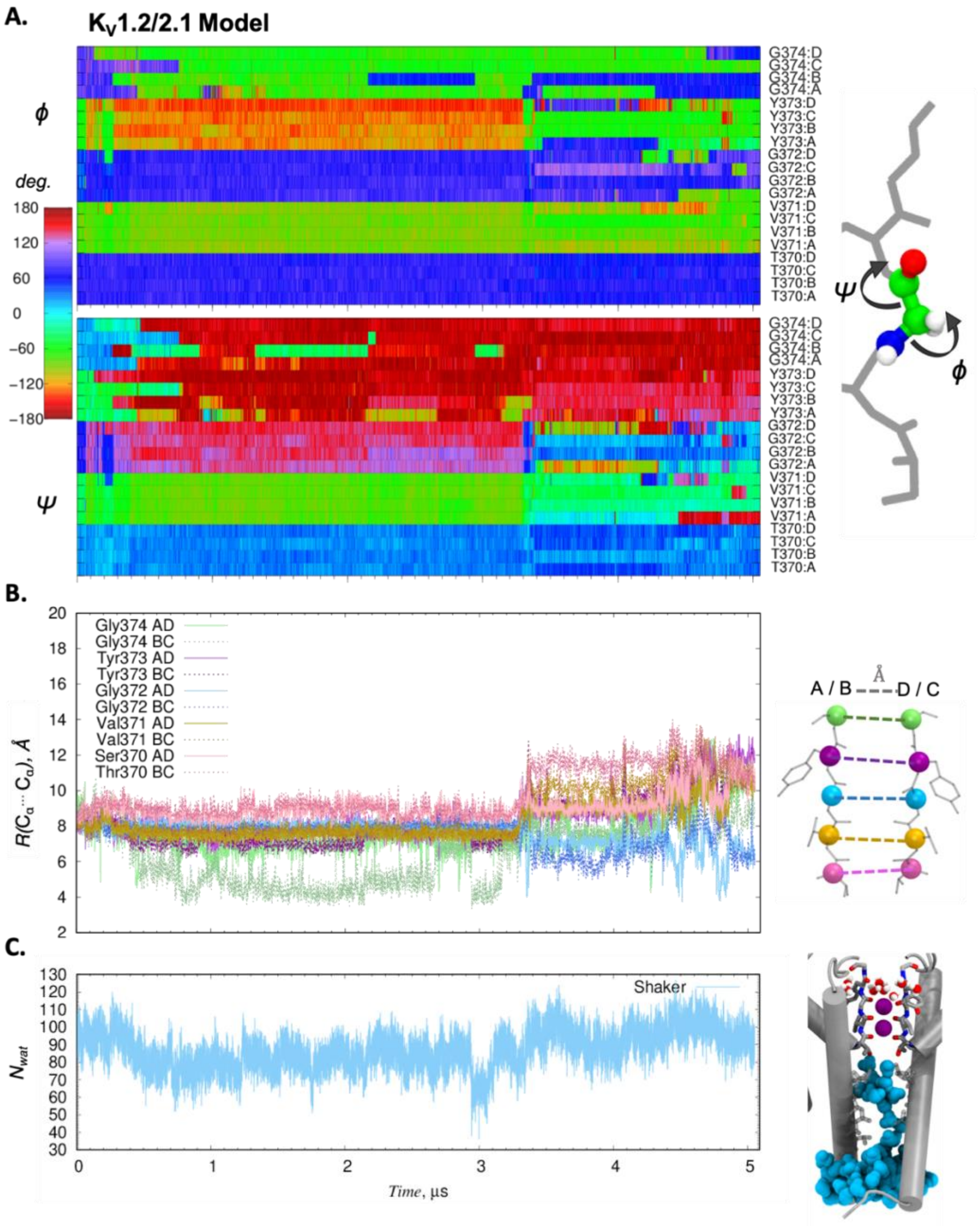

**Figure S8. Structural analysis of $K_V1.2/2.1$ chimera from MD simulations under applied 750 mV voltage. (A)** *Left:* Time series of $\varphi$ and $\psi$ protein backbone dihedral angles of SF residues TVGYG. *Right:* Graphical representation of $\varphi$ and $\psi$ dihedral angles in the SF. **(B)** *Left*: Time series of distances between protein backbone $C_\alpha$ atoms between SF residues in the opposing chains. *Right:* Depiction of the distances being measured, with protein chains "A" and "B" opposite to "D" and "C", respectively. **(C)** *Left:* Time series of pore hydration as measured by number of water molecules within the pore. *Right:* Snapshot the lowest degree of pore hydration ($t$ = 2.998 μs) as measured by water molecules within the pore. Relevant water molecules occupying pore are colored light-blue and correspond to the plotted quantity on the left. Water molecules behind SF are red/white.

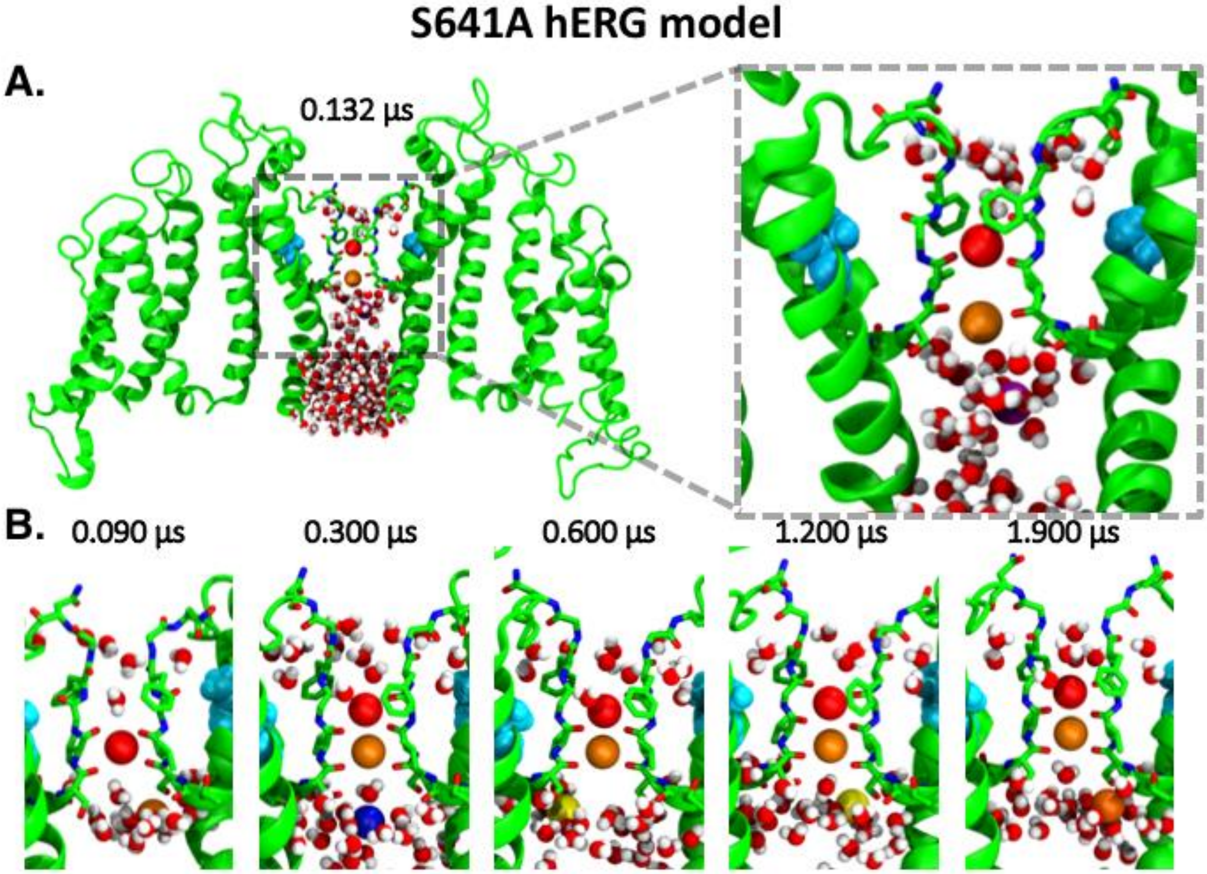

**Figure S9. $K^+$ pore positions for S641A hERG mutant channel model under an applied 750 mV voltage. (A)** Representative frame showing two opposite protein chains (green ribbons with SF S624-G628, S6 helix Y652 and F656 residues shown as sticks with red O and blue N, mutated Ala 641 residue shown as cyan in a space-filling representation), pore ions (colored spheres) and waters (red/white). Inset on the right shows closed-up view with SF pinched at the top clearly visible. ). **(B)** Close-up views of the channel SF in the same representation at different time points, showing no $K^+$ ion translocation for this hERG mutant during a 2 μs simulation with applied 750 mV voltage.

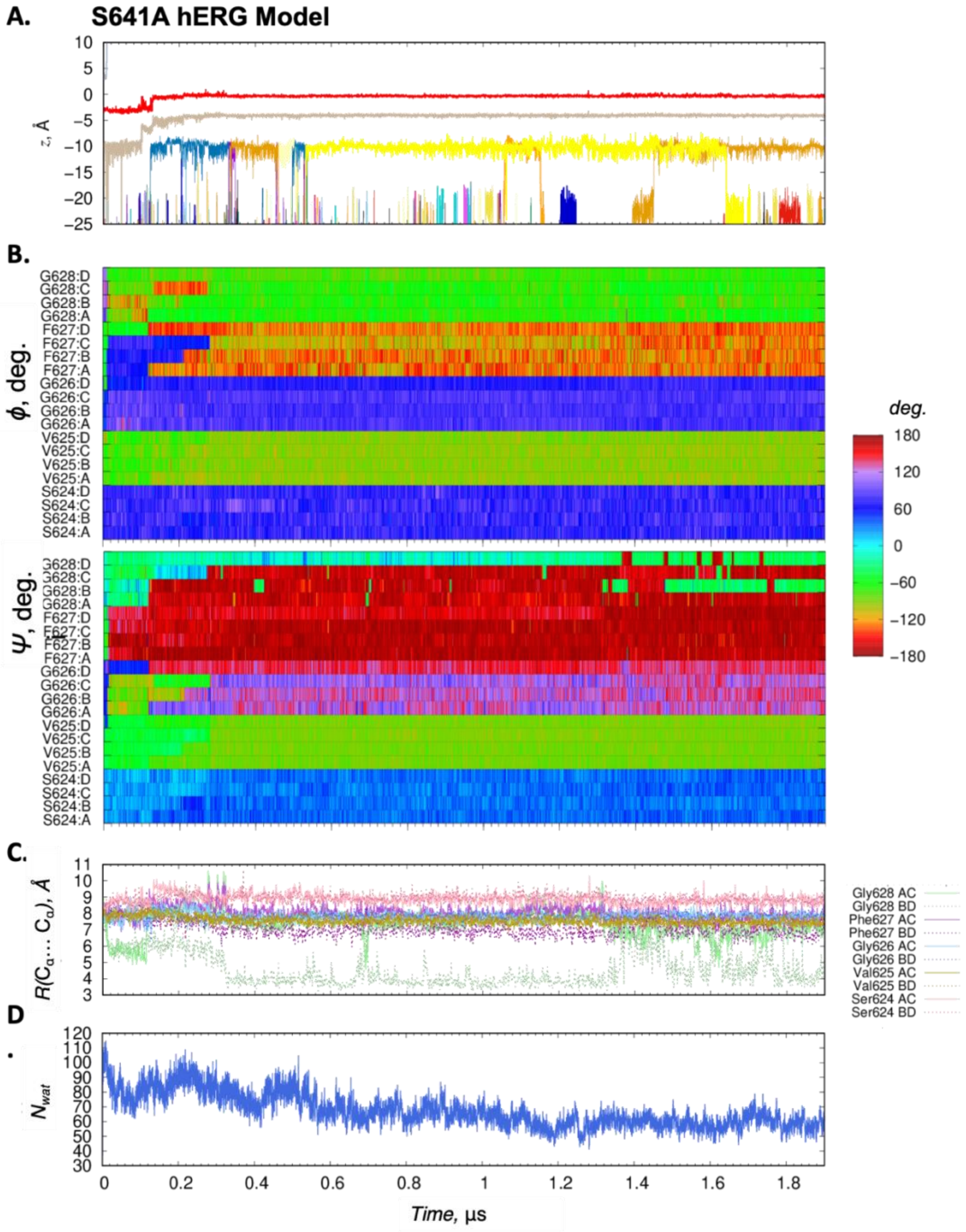

**Figure S10. Structural analysis of S641A hERG mutant model from MD simulations under applied 750 mV voltage. (A)** Time series depicting z-positions of $K^+$ ions (with respect to SF backbone center of mass) in the channel pore indicates no ion conduction in ~ 2 μs. **(B)** Time series of $\varphi$ and $\psi$ protein backbone dihedral angles of SF residues SVGFG for each chain. **(C)** Time series of distances between protein backbone $C_\alpha$ atoms between SF residues in the opposing chains ("AC" or "BD"). **(D)** Time series of pore hydration as measured by number of water molecules within the pore.

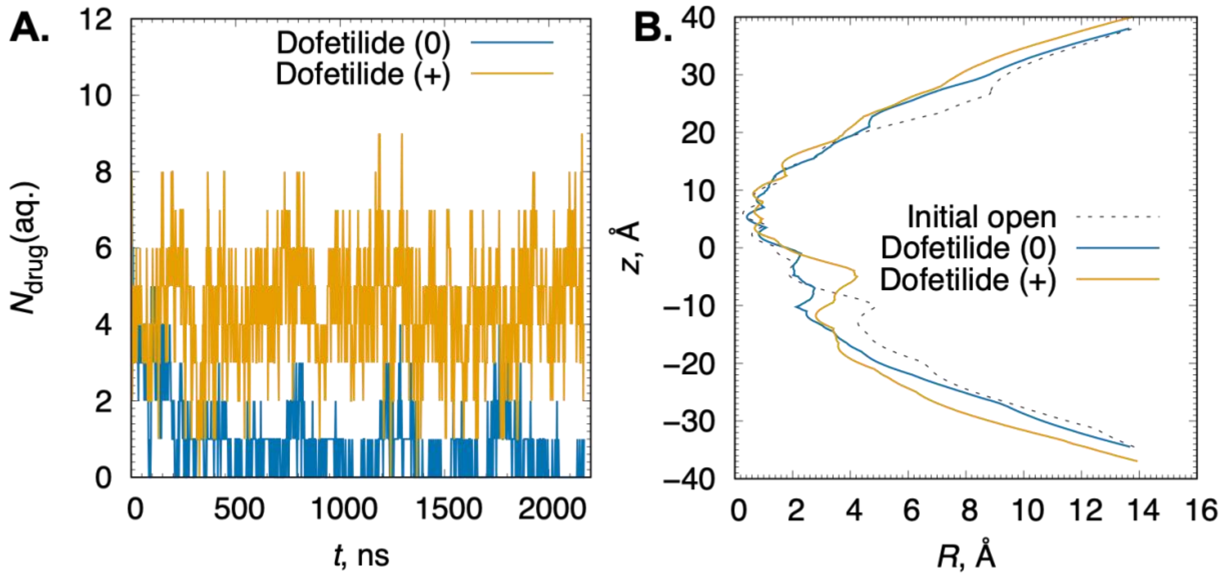

**Figure S11. Time series of number of dofetilide molecules in bulk aqueous solution (A) and channel pore radius profiles (B) from unbiased drug "flooding" hERG MD simulations. (A)** Number of drug molecules (Dofetilide (+) – orange and Dofetilide (0) – blue traces) in aqueous solution, $N_{\mathrm{drug}}$(aq.) over the course of the MD trajectory, with drug molecules considered membrane or protein bound when at least one of their non-hydrogen atoms was located within 3.5 Å from protein or lipid non-hydrogen atoms. **(B)** Comparison of pore radius profiles of hERG channel at the beginning (dashed black curve) of MD simulations, and at the end of ~2.5 μs trajectories of hERG MD simulations in the presence of Dofetilide (+) (solid orange curve) and Dofetilide (0) (blue solid curve).

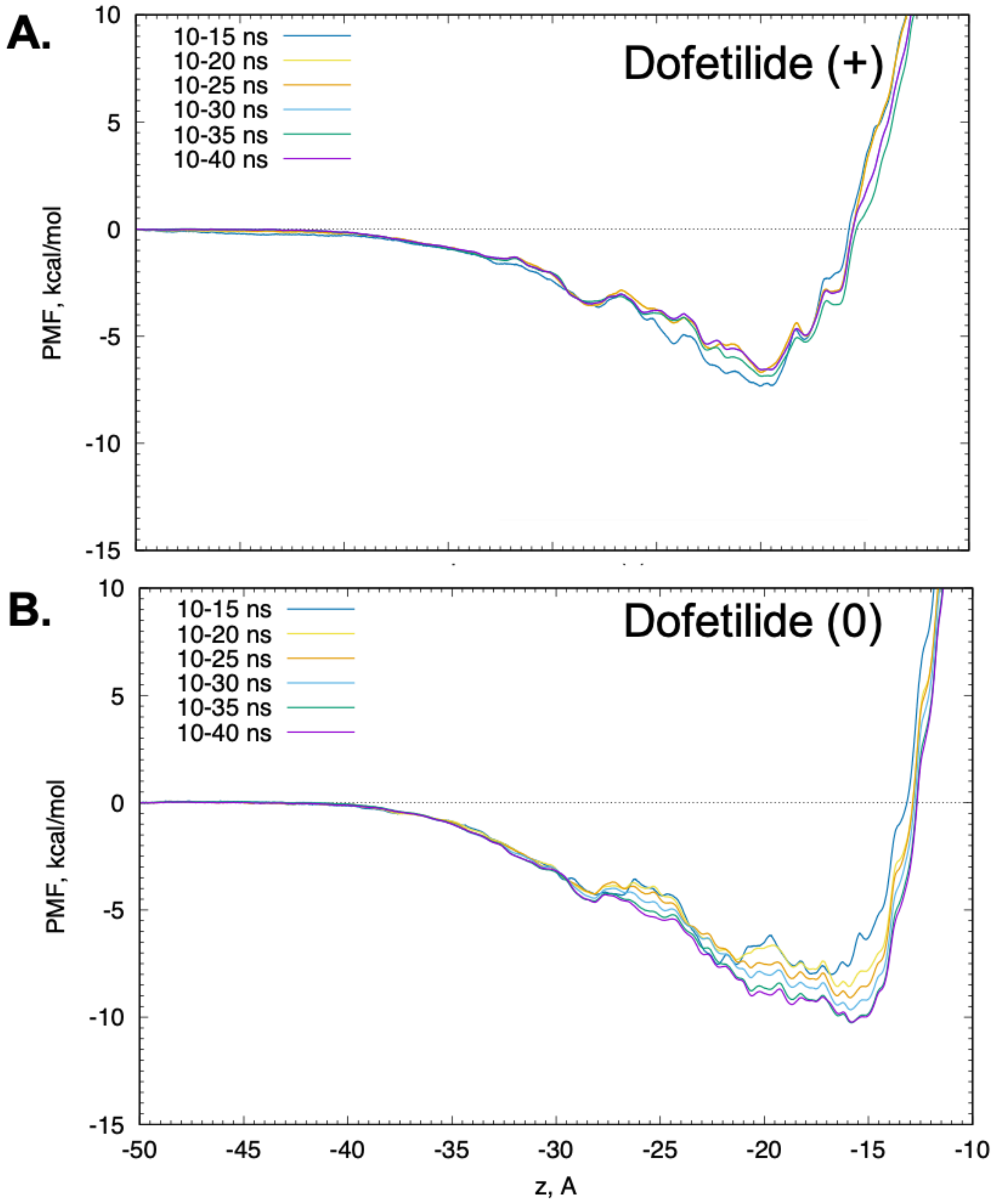

**Figure S12. Convergence of free energy or potential of mean force (PMF) profiles for drug binding to the hERG channel computed from umbrella sampling MD simulations.** **A.** Dofetilide (+) and **B.** Dofetilide (0) profiles computed using weighted histogram analysis method (WHAM) discarding first 10 ns for each umbrella sampling window, and incrementally including data from additional 5 ns blocks up to 40 ns.

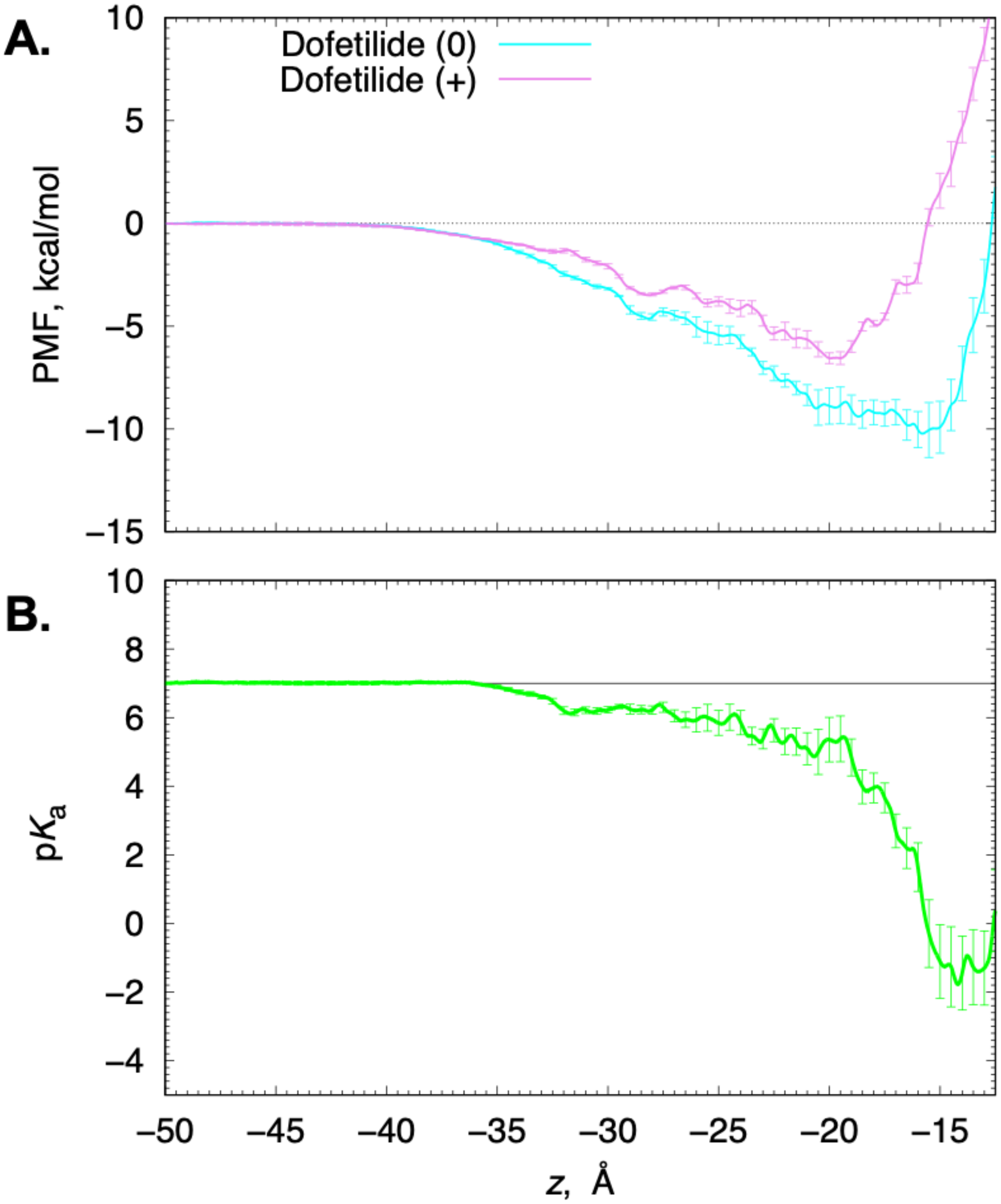

**Figure S13. Potential of mean force (PMF) and corresponding p$K_a$ profiles for dofetilide binding to hERG. (A)** PMF profiles of neutral (cyan) and charged (magenta) dofetilide binding to the hERG channel. **(B)** p$K_a$ profiles computed from PMFs in panel **A** (aqueous p$K_a$ = 7.0 is shown as a horizontal black line). Error bars are computed as standard errors from 5-ns block averaging.

#### SI Appendix

#### SI A1. Optimized force field topology and parameters for charged dofetilide

```
* Initial topologies generated by
* CHARMM General Force Field (CGenFF) program version 1.0.0
* For use with CGenFF version 3.0.1
36 1

! "penalty" is the highest penalty score of the associated parameters.
! Penalties lower than 10 indicate the analogy is fair; penalties between 10
! and 50 mean some basic validation is recommended; penalties higher than
! 50 indicate poor analogy and mandate extensive validation/optimization.

!=============================================================
! Dofetilide(+)
!=============================================================

RESI DOF1       1.000
GROUP            ! CHARGE   CH_PENALTY
ATOM C1     CG334   0.149
ATOM N1     NG3P1  -0.006
ATOM C2     CG324  -0.495
ATOM C3     CG321  -0.250
ATOM C4     CG2R61  0.310
ATOM C5     CG2R61 -0.115 !    8.929
ATOM C6     CG2R61 -0.111 !    0.000
ATOM C7     CG2R61  0.219 !    0.000
ATOM C8     CG2R61 -0.111 !    0.000
ATOM C9     CG2R61 -0.115 !    8.929
ATOM N2     NG311  -0.466 !    0.000
ATOM S1     SG3O2   0.439 !    0.000
ATOM O1     OG2P1  -0.384 !    0.000
ATOM O2     OG2P1  -0.384 !    0.000
ATOM C10    CG331  -0.019 !    0.000
ATOM C11    CG324   0.162 !    1.152
ATOM C12    CG321  -0.014 !    1.152
ATOM O3     OG301  -0.391 !    0.861
ATOM C13    CG2R61  0.219 !    0.850
ATOM C14    CG2R61 -0.113 !    0.000
ATOM C15    CG2R61 -0.115 !    0.000
ATOM C16    CG2R61  0.219 !    0.000
ATOM C17    CG2R61 -0.115 !    0.000
ATOM C18    CG2R61 -0.113 !    0.000
ATOM N3     NG311  -0.466 !    0.000
ATOM S2     SG3O2   0.439 !    0.000
ATOM O4     OG2P1  -0.384 !    0.000
ATOM O5     OG2P1  -0.384 !    0.000
ATOM C19    CG331  -0.019 !    0.000
ATOM H1     HGA3    0.090 !    0.000
ATOM H2     HGA3    0.090 !    0.000
ATOM H3     HGA3    0.090 !    0.000
ATOM H4     HGA2    0.090 !    2.455
ATOM H5     HGA2    0.090 !    2.455
ATOM H6     HGA2    0.090 !    0.000
ATOM H7     HGA2    0.090 !    0.000
ATOM H8     HGR61   0.115 !    0.000
ATOM H9     HGR61   0.115 !    0.000
ATOM H10    HGR61   0.115 !    0.000
ATOM H11    HGR61   0.115 !    0.000
ATOM H12    HGP1    0.323 !    0.000
ATOM H13    HGA3    0.090 !    0.000
ATOM H14    HGA3    0.090 !    0.000
ATOM H15    HGA3    0.090 !    0.000
ATOM H16    HGA2    0.090 !    0.075
ATOM H17    HGA2    0.090 !    0.075
ATOM H18    HGA2    0.090 !    0.000
ATOM H19    HGA2    0.090 !    0.000
ATOM H20    HGR61   0.115 !    0.000
```

```
ATOM H21    HGR61   0.115 !    0.000
ATOM H22    HGR61   0.115 !    0.000
ATOM H23    HGR61   0.115 !    0.000
ATOM H24    HGP1    0.323 !    0.000
ATOM H25    HGA3    0.090 !    0.000
ATOM H26    HGA3    0.090 !    0.000
ATOM H27    HGA3    0.090 !    0.000
ATOM HN     HGP2    0.318 !    0.000

BOND C1   N1
BOND C1   H1
BOND C1   H2
BOND C1   H3
BOND N1   C2
BOND N1   C11
BOND N1   HN
BOND C2   C3
BOND C2   H4
BOND C2   H5
BOND C3   C4
BOND C3   H6
BOND C3   H7
BOND C4   C9
BOND C4   C5
BOND C5   C6
BOND C5   H8
BOND C6   C7
BOND C6   H9
BOND C7   C8
BOND C7   N2
BOND C8   C9
BOND C8   H10
BOND C9   H11
BOND N2   S1
BOND N2   H12
BOND S1   O1
BOND S1   O2
BOND S1   C10
BOND C10  H13
BOND C10  H14
BOND C10  H15
BOND C11  C12
BOND C11  H16
BOND C11  H17
BOND C12  O3
BOND C12  H18
BOND C12  H19
BOND O3   C13
BOND C13  C18
BOND C13  C14
BOND C14  C15
BOND C14  H20
BOND C15  C16
BOND C15  H21
BOND C16  C17
BOND C16  N3
BOND C17  C18
BOND C17  H22
BOND C18  H23
BOND N3   S2
BOND N3   H24
BOND S2   O4
BOND S2   O5
BOND S2   C19
BOND C19  H25
BOND C19  H26
BOND C19  H27

* Initial parameters generated by analogy by
* CHARMM General Force Field (CGenFF) program version 1.0.0
* For use with CGenFF version 3.0.1
```

```
! Penalties lower than 10 indicate the analogy is fair; penalties between 10
! and 50 mean some basic validation is recommended; penalties higher than
! 50 indicate poor analogy and mandate extensive validation/optimization.

!=============================================================
! Dofetilide(+)
!=============================================================

BONDS

ANGLES
CG2R61 CG321  CG324    51.80    107.50 ! ZINC00 , from CG2R61 CG321 CG314, penalty= 0.6
CG324  CG321  OG301    75.70    110.10 ! ZINC00 , from CG324 CG321 OG302, penalty= 0.5

DIHEDRALS
CG2R61 CG2R61 CG321   CG324       0.2300   2    180.00 ! ZINC00 , from CG2R61 CG2R61 CG321 CG314,
penalty= 0.6
CG2R61 CG321   CG324   NG3P1       0.2000   3      0.00 ! ZINC00 , from NG3P3 CG314 CG321 CG2R61,
penalty= 5.5
CG2R61 CG321  CG324  HGA2      0.0400  3     0.00 ! ZINC00 , from CG2R61 CG321 CG321 HGA2, penalty=
1
OG301  CG321  CG324  NG3P1      3.3000  1   180.00 ! ZINC00 , from OG302 CG321 CG324 NG3P0, penalty=
1.7
OG301  CG321  CG324  NG3P1     -0.4000  3   180.00 ! ZINC00 , from OG302 CG321 CG324 NG3P0, penalty=
1.7
OG301  CG321  CG324  HGA2       0.1900  3     0.00 ! ZINC00 , from OG301 CG321 CG321 HGA2, penalty=
1
CG324  CG321  OG301  CG2R61     2.9990  1   180.00
CG324  CG321  OG301  CG2R61     2.4130  2   180.00
CG324  CG321  OG301  CG2R61     2.3380  3     0.00
```

#### SI A2. Optimized force field topology and parameters for neutral dofetilide

```
* Initial topologies generated by
* CHARMM General Force Field (CGenFF) program version 1.0.0
* For use with CGenFF version 3.0.1
36 1

! "penalty" is the highest penalty score of the associated parameters.
! Penalties lower than 10 indicate the analogy is fair; penalties between 10
! and 50 mean some basic validation is recommended; penalties higher than
! 50 indicate poor analogy and mandate extensive validation/optimization.

!=============================================================
! Dofetilide(0)
!=============================================================

RESI DOF0       0.000
GROUP            ! CHARGE   CH_PENALTY
ATOM C1     CG331  -0.392
ATOM N1     NG301  -0.401
ATOM C2     CG321   0.240
ATOM C3     CG321  -0.559
ATOM C4     CG2R61  0.296
ATOM C5     CG2R61 -0.115 !    0.000
ATOM C6     CG2R61 -0.111 !    0.000
ATOM C7     CG2R61  0.219 !    0.000
ATOM C8     CG2R61 -0.111 !    0.000
ATOM C9     CG2R61 -0.115 !    0.000
ATOM N2     NG311  -0.466 !    0.000
ATOM S1     SG3O2   0.439 !    0.000
ATOM O1     OG2P1  -0.384 !    0.000
ATOM O2     OG2P1  -0.384 !    0.000
ATOM C10    CG331  -0.019 !    0.000
ATOM C11    CG321   0.010
ATOM C12    CG321  -0.021 !    6.242
ATOM O3     OG301  -0.390 !    1.650
```

```
ATOM C13    CG2R61  0.219 !    0.000
ATOM C14    CG2R61 -0.113 !    0.000
ATOM C15    CG2R61 -0.115 !    0.000
ATOM C16    CG2R61  0.219 !    0.000
ATOM C17    CG2R61 -0.115 !    0.000
ATOM C18    CG2R61 -0.113 !    0.000
ATOM N3     NG311  -0.466 !    0.000
ATOM S2     SG3O2   0.439 !    0.000
ATOM O4     OG2P1  -0.384 !    0.000
ATOM O5     OG2P1  -0.384 !    0.000
ATOM C19    CG331  -0.019 !    0.000
ATOM H1     HGA3    0.090 !    3.536
ATOM H2     HGA3    0.090 !    3.536
ATOM H3     HGA3    0.090 !    3.536
ATOM H4     HGA2    0.090 !    3.536
ATOM H5     HGA2    0.090 !    3.536
ATOM H6     HGA2    0.090 !    0.480
ATOM H7     HGA2    0.090 !    0.480
ATOM H8     HGR61   0.115 !    0.000
ATOM H9     HGR61   0.115 !    0.000
ATOM H10    HGR61   0.115 !    0.000
ATOM H11    HGR61   0.115 !    0.000
ATOM H12    HGP1    0.323 !    0.000
ATOM H13    HGA3    0.090 !    0.000
ATOM H14    HGA3    0.090 !    0.000
ATOM H15    HGA3    0.090 !    0.000
ATOM H16    HGA2    0.090 !    3.536
ATOM H17    HGA2    0.090 !    3.536
ATOM H18    HGA2    0.090 !    0.480
ATOM H19    HGA2    0.090 !    0.480
ATOM H20    HGR61   0.115 !    0.000
ATOM H21    HGR61   0.115 !    0.000
ATOM H22    HGR61   0.115 !    0.000
ATOM H23    HGR61   0.115 !    0.000
ATOM H24    HGP1    0.323 !    0.000
ATOM H25    HGA3    0.090 !    0.000
ATOM H26    HGA3    0.090 !    0.000
ATOM H27    HGA3    0.090 !    0.000

BOND C1   N1
BOND C1   H1
BOND C1   H2
BOND C1   H3
BOND N1   C2
BOND N1   C11
BOND C2   C3
BOND C2   H4
BOND C2   H5
BOND C3   C4
BOND C3   H6
BOND C3   H7
BOND C4   C9
BOND C4   C5
BOND C5   C6
BOND C5   H8
BOND C6   C7
BOND C6   H9
BOND C7   C8
BOND C7   N2
BOND C8   C9
BOND C8   H10
BOND C9   H11
BOND N2   S1
BOND N2   H12
BOND S1   O1
BOND S1   O2
BOND S1   C10
BOND C10  H13
BOND C10  H14
BOND C10  H15
BOND C11  C12
```

```
BOND C11  H16
BOND C11  H17
BOND C12  O3
BOND C12  H18
BOND C12  H19
BOND O3   C13
BOND C13  C18
BOND C13  C14
BOND C14  C15
BOND C14  H20
BOND C15  C16
BOND C15  H21
BOND C16  C17
BOND C16  N3
BOND C17  C18
BOND C17  H22
BOND C18  H23
BOND N3   S2
BOND N3   H24
BOND S2   O4
BOND S2   O5
BOND S2   C19
BOND C19  H25
BOND C19  H26
BOND C19  H27

* Initial parameters generated by analogy by
* CHARMM General Force Field (CGenFF) program version 1.0.0
* For use with CGenFF version 3.0.1

! Penalties lower than 10 indicate the analogy is fair; penalties between 10
! and 50 mean some basic validation is recommended; penalties higher than
! 50 indicate poor analogy and mandate extensive validation/optimization.

BONDS
CG321  NG301   263.00     1.4740 ! ZINC00 , from CG321 NG311, penalty= 5
CG331  NG301   255.00     1.4630 ! ZINC00 , from CG331 NG311, penalty= 5

ANGLES
CG321  CG321  NG301    43.70    112.20 ! ZINC00 , from CG331 CG321 NG311, penalty= 1.5
NG301  CG321  HGA2     32.40    109.50   50.00   2.13000 ! ZINC00 , from NG311 CG321 HGA2, penalty=
0.6
NG301  CG331  HGA3     30.50    109.70   50.00   2.14000 ! ZINC00 , from NG311 CG331 HGA3, penalty=
0.6
CG321  NG301  CG321    42.452   104.717
CG321  NG301  CG331    59.281   106.264

DIHEDRALS
CG2R61 CG321  CG321  NG301      0.3650  3     0.00
NG301  CG321  CG321  OG301      3.0000  1   180.00
NG301  CG321  CG321  OG301      0.9850  2     0.00
NG301  CG321  CG321  HGA2       0.1600  3     0.00 ! ZINC00 , from NG311 CG321 CG331 HGA3, penalty=
6.6
CG321  CG321  NG301  CG321      1.7910  1     0.00
CG321  CG321  NG301  CG321      0.7430  2     0.00
CG321  CG321  NG301  CG321      0.1570  3     0.00
CG321  CG321  NG301  CG331      0.5940  1     0.00
CG321  CG321  NG301  CG331      0.1350  2     0.00
CG321  CG321  NG301  CG331      1.0280  3     0.00
HGA2   CG321  NG301  CG321      0.3570  3   180.00
HGA2   CG321  NG301  CG331      0.7040  3     0.00
HGA3   CG331  NG301  CG321      0.8520  3   180.00
```
